# Global disinhibition and corticospinal plasticity for drastic recovery after spinal cord injury

**DOI:** 10.1101/2023.03.15.532773

**Authors:** Reona Yamaguchi, Satoko Ueno, Toshinari Kawasaki, Zenas C. Chao, Masahiro Mitsuhashi, Kaoru Isa, Tomohiko Takei, Kenta Kobayashi, Jun Takahashi, Hirotaka Onoe, Tadashi Isa

## Abstract

The induction of large-scale plasticity in the adult brain should be key for recovery from severe damage of the central nervous system. Here, drastic motor recovery was observed after subhemisection spinal cord injury in macaques that received intensive training and cortical electrical stimulation. During recovery, movement-related activity increased in ipsilesional sensorimotor areas and functional connectivity from ipsilesional to contralesional areas was strengthened. Electrical stimulation applied widely across bilateral sensorimotor areas induced muscle twitches in affected and intact forelimbs. The interhemispheric inhibition observed before injury was switched to facilitation. Furthermore, massive re-routing occurred in corticospinal axons from the contralesional motor cortex. Such global disinhibition and massive plasticity would open the workspace for the reorganization of motor networks to recruit novel areas for recovery.

**One Sentence Summary:** Global disinhibition and corticospinal plasticity for drastic recovery after spinal cord injury in macaque monkeys.

## Main Text

The difficulty in curing from severe damage of the central nervous system is due to the limited plasticity of adult brains. However, recent studies revealed massive plasticity during recovery from neuronal injuries, where a variety of brain areas, which are not normally involved in motor control, become activated and contribute to the direct control of movements. For instance, many neuroimaging studies have reported that the activity of motor/premotor cortices ipsilateral to the affected hand is increased in patients with spinal cord injury or stroke (*1–3*). In a nonhuman primate model of partial spinal cord injury targeting the corticospinal tract, we demonstrated that the ipsilesional motor cortex became involved in the control of grasping movements during the early recovery stage, while the premotor cortex contributes directly to hand movements during the late recovery stage (*4, 5*). The remote effect of a particular neuronal injury to brain areas that are directly or indirectly connected with it is classically called diaschisis (*6*), and recently, this concept has recaptured attention with its possible role in the recovery process after neuronal injury (*7–10*). However, its underlying neuronal mechanisms are elusive. Here, as possible mechanisms supporting diaschisis for the postlesion reorganization of large-scale brain networks, we propose “global disinhibition” and “massive plasticity”, which would release the neuronal workspace for re-learning of motor control strategies and recruit novel areas for recovery.

Two male macaque monkeys (*Macaca fuscata*, monkeys M and H, 6.5 and 5.2 kg bodyweight, respectively) were trained to sit on a monkey chair and perform a reach-and-grasp task to retrieve a small piece of sweet potato through a narrow slit and eat it (Fig. 1A). Electromyography (EMG) electrodes were chronically implanted subcutaneously in multiple muscles (monkey M: 12 muscles; monkey H: 13 muscles) spanning from the shoulder to the digits. Additionally, 36- and 56-channel electrocorticography (ECoG) electrodes were implanted on both sides of the premotor (PM), primary motor (M1), and primary somatosensory (S1) areas (Figs. 1B and S1) in monkeys M and H, respectively, for chronic recording of neuronal activity (*5, 11*), and the effect of electrical stimulation of the cortical surface in inducing muscle twitches was tested. The kinematics of reaching and grasping movements were analyzed by a motion capture system with markers on their hand. We conducted subhemisection spinal cord injury at the border between the C4 and C5 segments; the lesion was larger in monkey M than in monkey H. During surgery, we confirmed the completeness of the injury as shown in the video-captured images in Fig. S3, but, as shown in Figs. 1D and S2, some “islands” of partly intact areas could be observed (gray-hatched areas) in which many glial cells were distributed and Klüver-Barrera staining was faint, surrounded by the glial scar (black areas). Our interpretation of these zones is provided at the end of the Discussion.

**Fig. 1.**
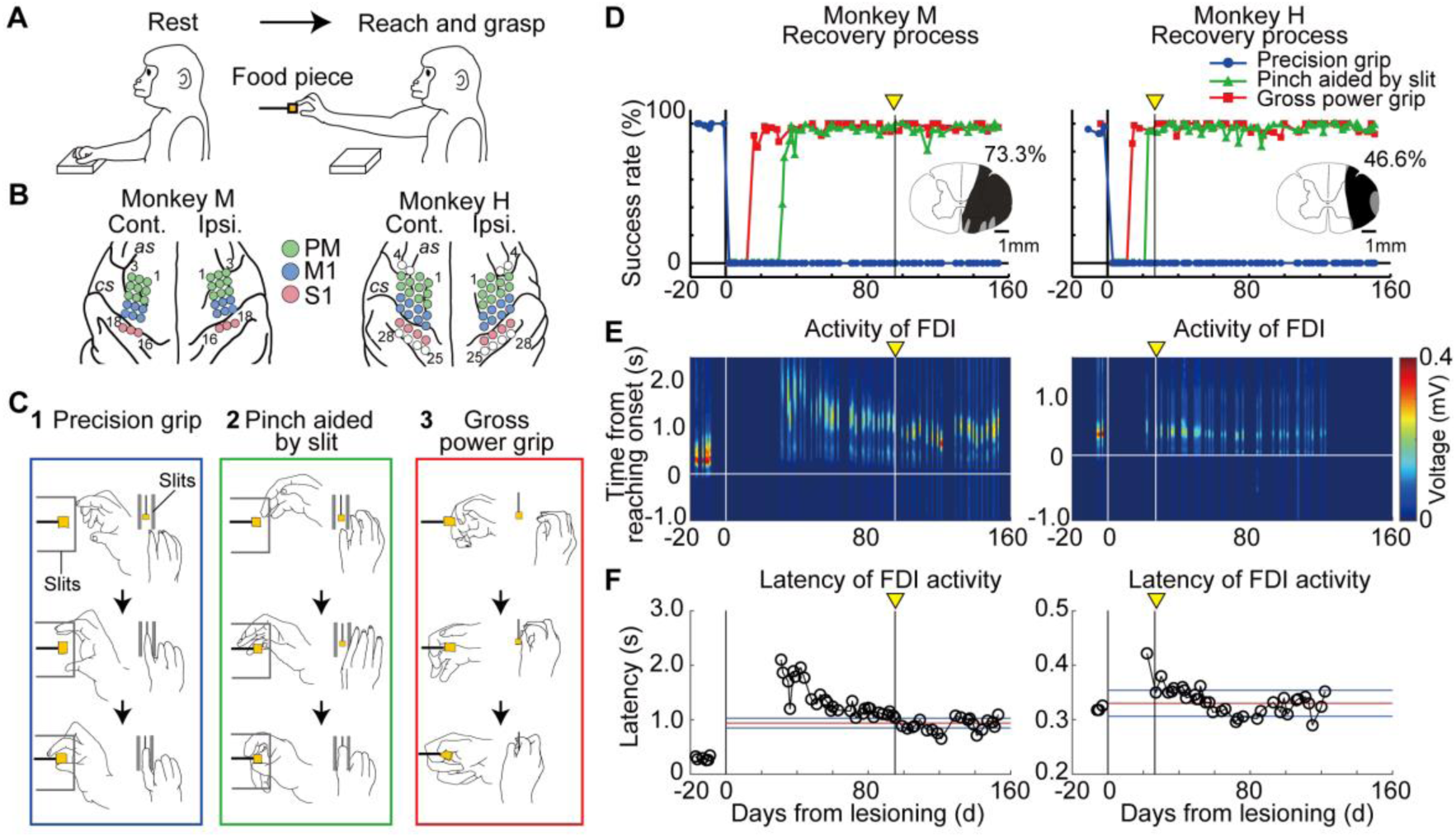
Recovery process of hand movements after subhemisection. (A) Reach-and-grasp task. (B) Arrangement of the electrocorticography (ECoG) electrodes. as, arcuate sulcus; cs, central sulcus. (C) Drawings of the three grasping types from lateral and top views. A food piece was presented in the slit. 1) Precision grip; 2) Pinch aided by the slit; 3) Gross power grip (in the absence of the slit). (D) Recovery curves. Time course of the success ratio of the three grasping types. The black area in the inset indicates the site of spinal cord injury. The percentage value beside the spinal cord illustration shows the proportion of tissue that was lesioned. (E) Time course of first dorsal interosseous (FDI) activity during the task. (F) Time course of peak FDI activity. Yellow arrowheads indicate the time when the recovery of FDI activity was saturated after lesioning. Red and blue horizontal lines indicate the mean and standard deviation for the last 10 days, respectively.

### Recovery after subhemisection of the cervical spinal cord

After subhemisection, both monkeys demonstrated severe paralysis of the affected forelimb (Fig. 1D, Supplementary Movies 1 and 2); however, through intensive training, their reaching and grasping movements recovered considerably. Although precision grip did not recover as the prelesion state (Fig. 1C1 and blue in Fig. 1D), first, coarse power grip without the slit (Fig. 1C3 and red in Fig. 1D) and then pinching aided by the slit edge (Fig. 1C2 and green in Fig. 1D) recovered in several weeks. In addition, the amplitude of EMG activity of the first dorsal interosseous (FDI) did not recover to the prelesion level (Fig. 1E), and recovery for peak latency of FDI activity was saturated on Days 95 and 27 in monkeys M and H, respectively (vertical lines in Fig. 1E, F). The recovery in these monkeys was surprisingly faster than in previous reports of hemisection or subhemisection models (*12*-*14*).

### Change in the activity and signal flow through the cortical networks

Time frequency analysis of ECoG activity of the ipsilesional PM and M1 showed it was increased during movement at α (8–12 Hz) band throughout recovery process (Figs. 2B and S4–8). In addition, activity of contralesional PM and M1 increased at α band after lesioning (Fig. 2A). The activity of the high-γ band (71–120 Hz) peaked during the late phase of recovery process in monkey M, which showed slower recovery, while it peaked earlier in monkey H, which showed faster recovery (Figs. S5–8). The connectivity structure, as assessed by Granger causality, primarily comprised of signal flow from the ipsilesional PM to contralesional PM/M1/S1, was enhanced at the α band during recovery in both monkeys (Figs. 2C–G and S9–14). Thus, the activity of the ipsilesional PM and signal flow from the ipsilesional PM were markedly enhanced during recovery.

**Fig. 2.**
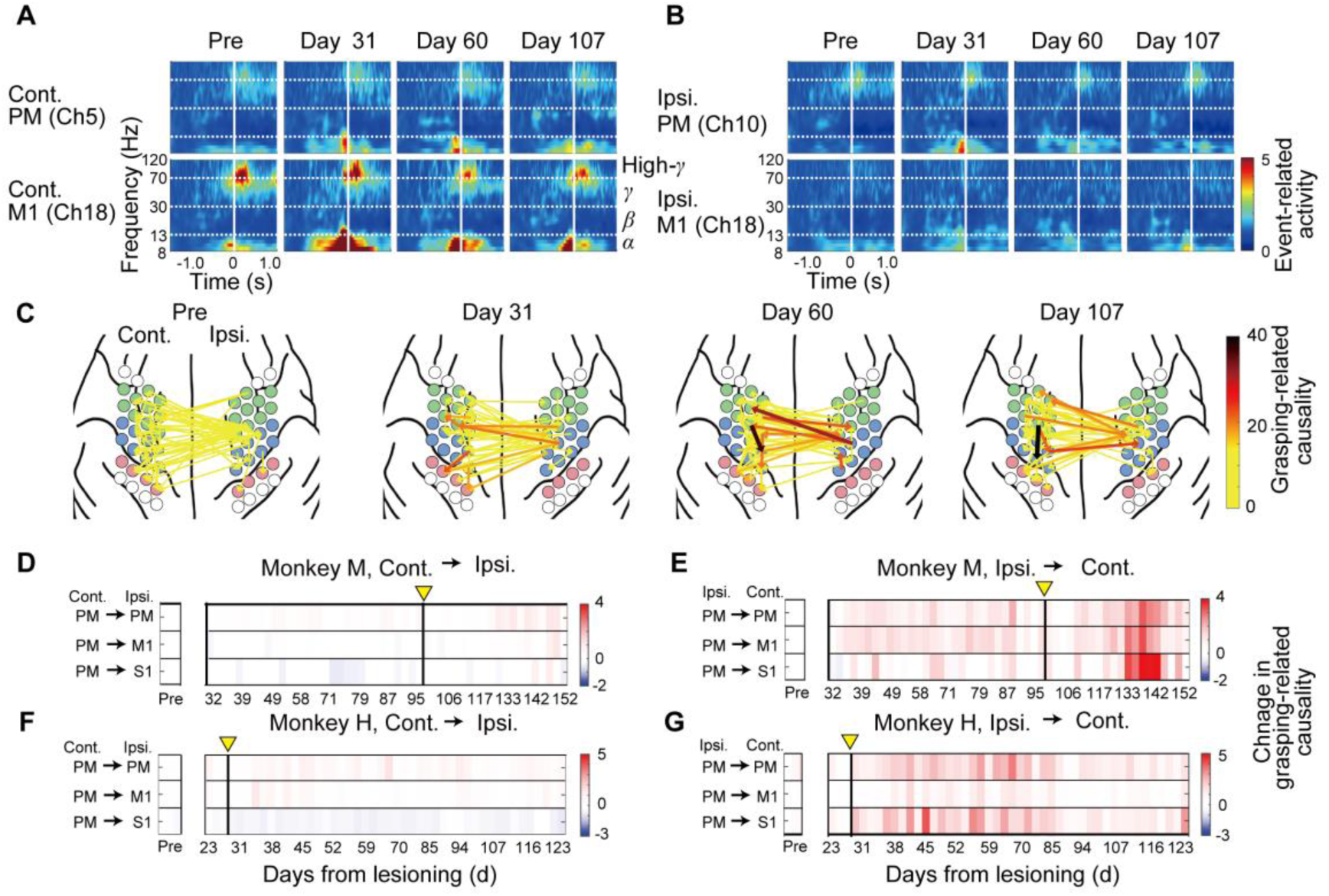
Dynamics of cortical activation and interaction. (A) Example of movement-related activity in the contralesional primary motor (M1) and premotor (PM) areas before and after lesioning for monkey H. Ch, channel. (B) Example of movement-related activity in ipsilesional M1and PM before and after lesioning for monkey H. (C) Example of grasping-related connectivity in the α band for monkey H. The top 5 % connections are shown. (D) Dynamics of grasping-related connectivity from the contralesional PM to ipsilesional PM, M1, and primary somatosensory (S1) areas in the α band for monkey M. Color indicates activity change in grasping-related connectivity compared with activity in the preoperative stage. Blue and red areas indicate statistically significant deactivated and activated components, respectively. (E) Dynamics of grasping-related connectivity from the ipsilesional PM to contralesional PM, M1 and S1 for monkey M. The arrangement is the same as in (D). (F, G) Results for monkey H. The arrangements are the same as in (D) and (E), respectively.

### Cortical mapping with electrical stimulation

We then mapped the effects of electrical stimulation of the bilateral PM, M1 and S1 through each channel of the ECoG electrodes on both sides with a short train stimulus (3 mA, cathode, 0.5 ms duration, × 3 at 20 Hz) by observing muscle twitches in both forelimbs (Fig. 3A). The magnitude of muscle twitches was classified into four ranks (3 (dark blue) = muscle twitches with joint movements, 2 (blue) = visible muscle twitches, 1 (light blue) = invisible muscle twitches, 0 (white) = no response). Stimulation of the contralesional (left) M1 (channel [Ch] 11) induced muscle twitches in the shoulder, elbow, and fingers before lesioning (Fig. 3B, upper panels). After subhemisection, twitches of the fingers disappeared but rank 2 twitches (visible muscle twitches) were observed in the elbow and upper arm by Day 20 postoperatively. Then, the twitches expanded to the forearm, wrist and fingers on Day 24 and continued until Day 150. The extent and magnitude of twitches were gradually reduced after Day 108. Although the magnitude of twitches was smaller, a similar trend was observed in monkey H (Fig. 3B). Lower panels of Fig. 3B show the effect of stimulation through an ECoG channel on the ipsilesional PM (Ch 8) in monkey M. In this case, no twitches were induced before lesioning. However, they emerged in the proximal muscles on Days 21–136 and extended to the finger muscles on Days 60–67, and then their extent was reduced. Although the twitches were limited to the proximal muscles, a similar trend was observed in monkey H.

**Fig. 3.**
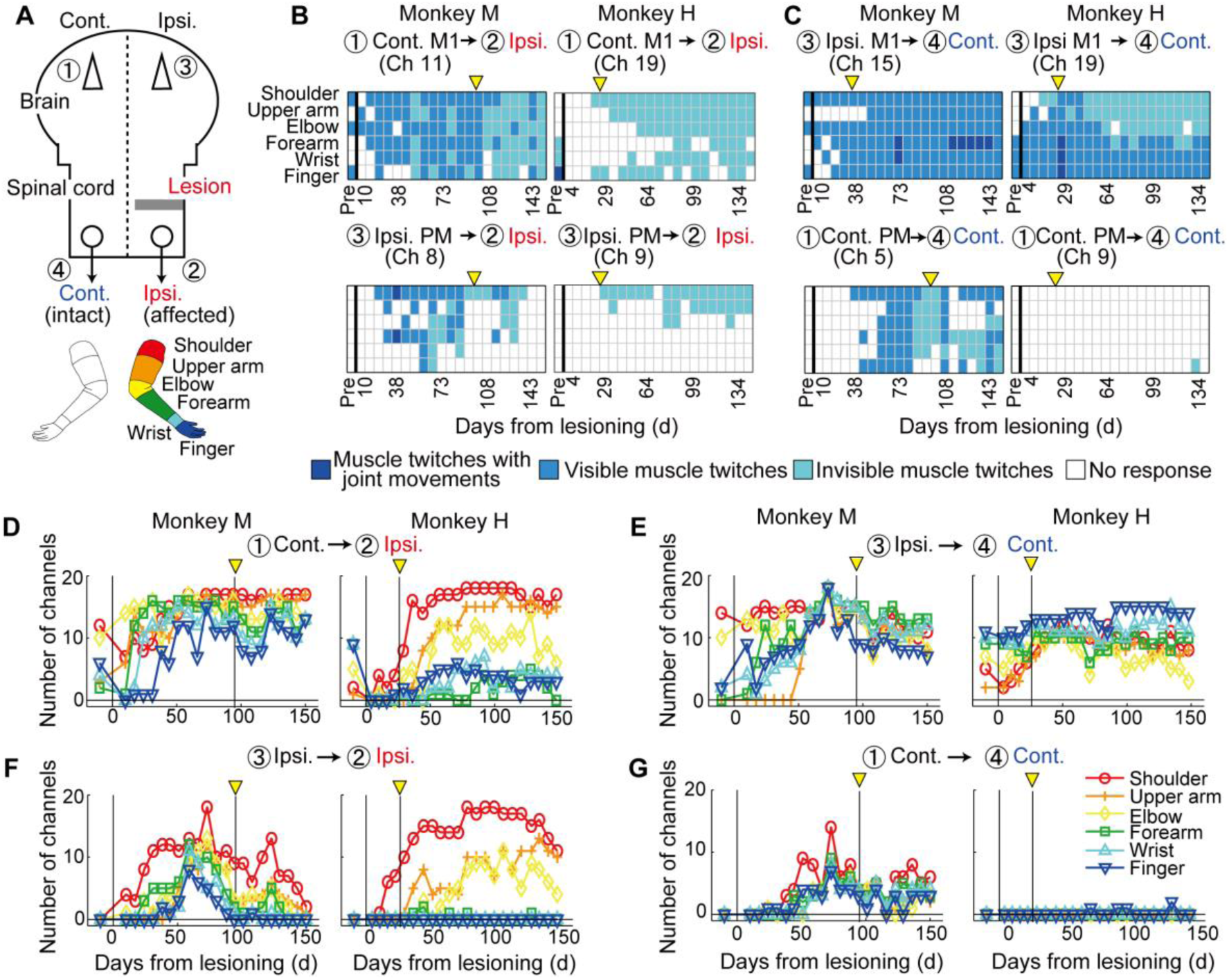
Effect of electrical stimulation. (A) Schematic showing the stimulation side and side of the body where muscle twitches were observed. Lower part shows the classification of body parts in which muscle twitches were observed. (B) Time course of induced muscle twitches on the affected side (right forelimb) by stimulation of the contralesional (left hemisphere) and ipsilesional (right hemisphere) sides. The magnitude of muscle twitches was classified into 4 ranks (3 (dark blue) = muscle twitches with joint movements, 2 (blue) = visible muscle twitches, 1 (light blue) = invisible muscle twitches, 0 (white) = no response). Ch, channel. (C) Time course of induced muscle twitches on the intact side (left forelimb) by stimulation of the ipsilesional (right hemisphere) and contralesional (left hemisphere) sides. (D) Change in the number of channels that induced muscle twitches in each body part on the affected side by stimulation of the contralesional side before and after lesioning. The total number of electrodes was 18. The magnitude of muscle twitches was ignored. (F) Results for stimulation of the ipsilesional side. (E) Change in the number of channels that induced muscle twitches in each body part on the intact side by stimulation of the ipsilesional side before and after lesioning. (G) Results for stimulation of the contralesional side.

The number of ECoG channels across the contralesional and ipsilesional PM, M1, and S1 that induced twitches in different muscle types is plotted in Figs. 3D, F, and S14-17, showing that wider areas spanning the PM, M1 and S1 on the contralesional and ipsilesional sides induced twitches in a wide extent of muscles from proximal to distal in the ipsilesional forelimb. More interestingly, such an expansion of the cortical areas whose stimulation induced muscle twitches was also observed in the contralesional (intact) forelimb muscles (Figs. 3C, E, G, and S18–21). The number of ECoG channels that induced muscle twitches increased not only in the ipsilesional cortical areas, which are primary movers (Fig. 3D), but also in the contralesional cortical areas, which are ipsilateral to the non-prime mover hand. This trend was clear in monkey M, and less clear in monkey H, although it could also be observed in a few channels.

### Switch from interhemispheric inhibition to interhemispheric facilitation

Thus, the PM, M1 and S1 on both sides appeared to have become highly excitable in terms of the induction of muscle twitches regardless of the target hand, both on the affected and intact sides. We hypothesized that the excitation/inhibition (E/I) balance might have shifted more to excitation in the global cortical circuits during the recovery process by way of disinhibition. In line with this hypothesis, we tested interhemispheric interactions using a conditioning-test paradigm. First, the amplitude of the evoked potentials in the contralesional M1 by stimulation of the ipsilesional PM was enhanced after lesioning (Figs. 4A, B, and S22). Furthermore, as shown in Fig. 4C, D (data from monkey H) and Figs. S23 and S24, the EMG responses to stimulation of the contralesional M1 (Ch18) recorded in the intrinsic hand muscles were used as the test responses and the effect of conditioning stimulus applied to the ipsilesional PM with a time intervals of 0–50 ms was examined. The effect of the conditioning stimulus was inhibitory before lesioning as reported in human studies (*15*); however, it became facilitatory during the early stage of recovery (Day 56).

**Fig. 4.**
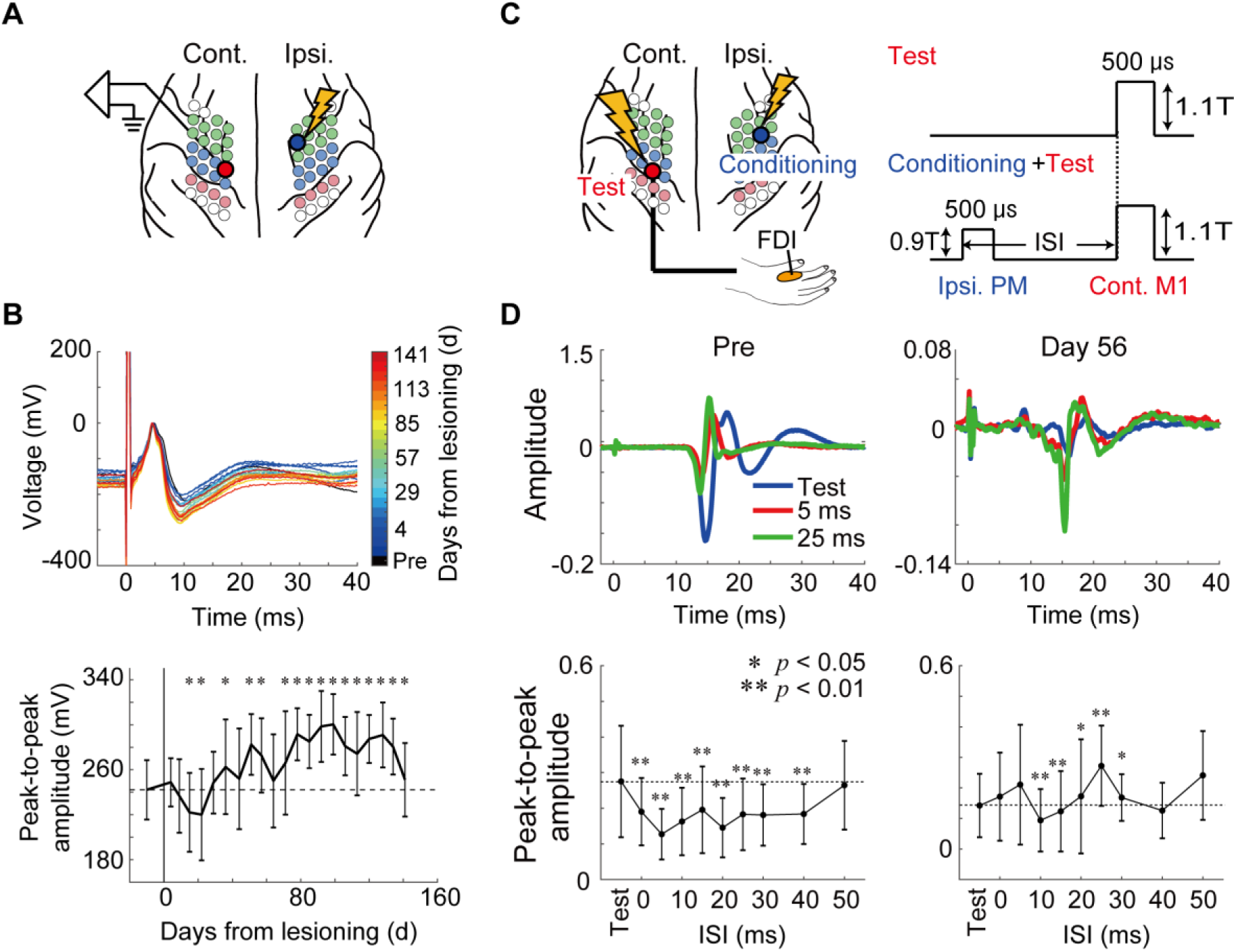
Interhemispheric interactions before and after spinal cord injury in monkey H. (A) Schematic of stimulation side (ipsilesional premotor [PM] area, channel [Ch] 5 in Fig. S1) and recording of the evoked potentials on the opposite side (contralesional primary motor [M1] area, Ch13 in Fig. S1) to the stimulation side. (B) Change in the shape of the evoked potentials and peak-to-peak amplitudes. In the bottom panel, the dotted line indicates the averaged peak-to-peak amplitude in the preoperative stage. The asterisks indicate a significant difference in peak-to-peak amplitude between experimental days and preoperative stage (p < 0.05, Wilcoxon rank-sum test). (C) Schematic drawing of the conditioning-test paradigm. The test stimulus was applied to the contralesional (left) M1 (Ch18 in Fig. S1) and the conditioning stimulus was given to the ipsilesional (right) PM (Ch11 in Fig. S1). The electromyography (EMG) responses were recorded from the right first dorsal interosseous (FDI). (D) Results of the conditioning test before (left) and at Day 56 after spinal cord injury (right). The upper panels show the superimposed records of the EMG responses with the test stimulus alone (blue) and with the conditioned stimulus (interstimulus interval [ISI] 5 ms, red; 25 ms, green). The lower panels indicate the averaged peak-to-peak responses with different ISIs. The horizontal dotted line indicates the magnitude of the test response alone. Asterisks indicate a significant difference in peak-to-peak amplitude between the conditioned and test responses (* p < 0.05, ** p < 0.01, Wilcoxon rank-sum test).

### Massive plasticity of corticospinal axons

The above results showed that interhemispheric inhibition switched to interhemispheric facilitation during the recovery process. However, one may wonder how the motor commands can reach hand motoneurons in these animals, despite the disruption of the spinal cord by lesioning in the downstream. Previous studies on a similar subhemisection macaque model suggested that the extension of corticospinal axons from the contralesional M1 to bypass the lesion (*14*), or from the axons of the uncrossed corticospinal tract (CST) that crossed the midline at lower cervical segments (*13*) might be involved in recovery (*16*). To clarify the neural circuit supporting recovery, we injected two different viral tracers in the bilateral M1 in the rostral bank of the central sulcus (Figs. 5A and S25A). Figs. 5B and S25B show the distribution of labeled axons at the pyramidal decussation and the C3 and C6 segments in monkeys M and H, respectively. The distribution of labeled axons originating from the ipsilesional M1 was almost the same as in the intact monkeys (left row in Figs. 5B and S25B). However, a substantial number of labeled axons originating from the contralesional M1 were distributed on the contralesional side of the pyramidal decussation, which descended into the contralesional side of the spinal cord and then crossed the midline to terminate in the ipsilesional gray matter at C6 (Fig. S27). To quantify the number of labeled axons, we counted them using a camera lucida in the gray and white matter between the contralesional and ipsilesional sides at each segment (Figs. 5C and S25C, Table S1). We calculated the proportion of labeled axons between the contralesional and ipsilesional sides at each segment in the regions of interest shown in Fig. S26. In the case of labeled axons originating from the ipsilesional M1, approximately 90% were distributed on the contralesional side at each segment as shown previously (*17, 18*). Conversely, the proportion of labeled axons originating from the contralesional M1 at the pyramidal decussation was 15.4% and 31.4% in monkeys M and H, respectively. Figs. 5D and S25D show the distribution of labeled axons in the Rexed laminae. We found that the labeled axons originating from the contralesional M1 terminated in laminae VI–IX on the ipsilesional side, which were the original targets before injury. These results suggested that the corticospinal axons originating from the contralesional M1 were re-routed in the pyramidal decussation, reconnected to the motoneurons below the lesion, and contributed to recovery.

**Fig. 5.**
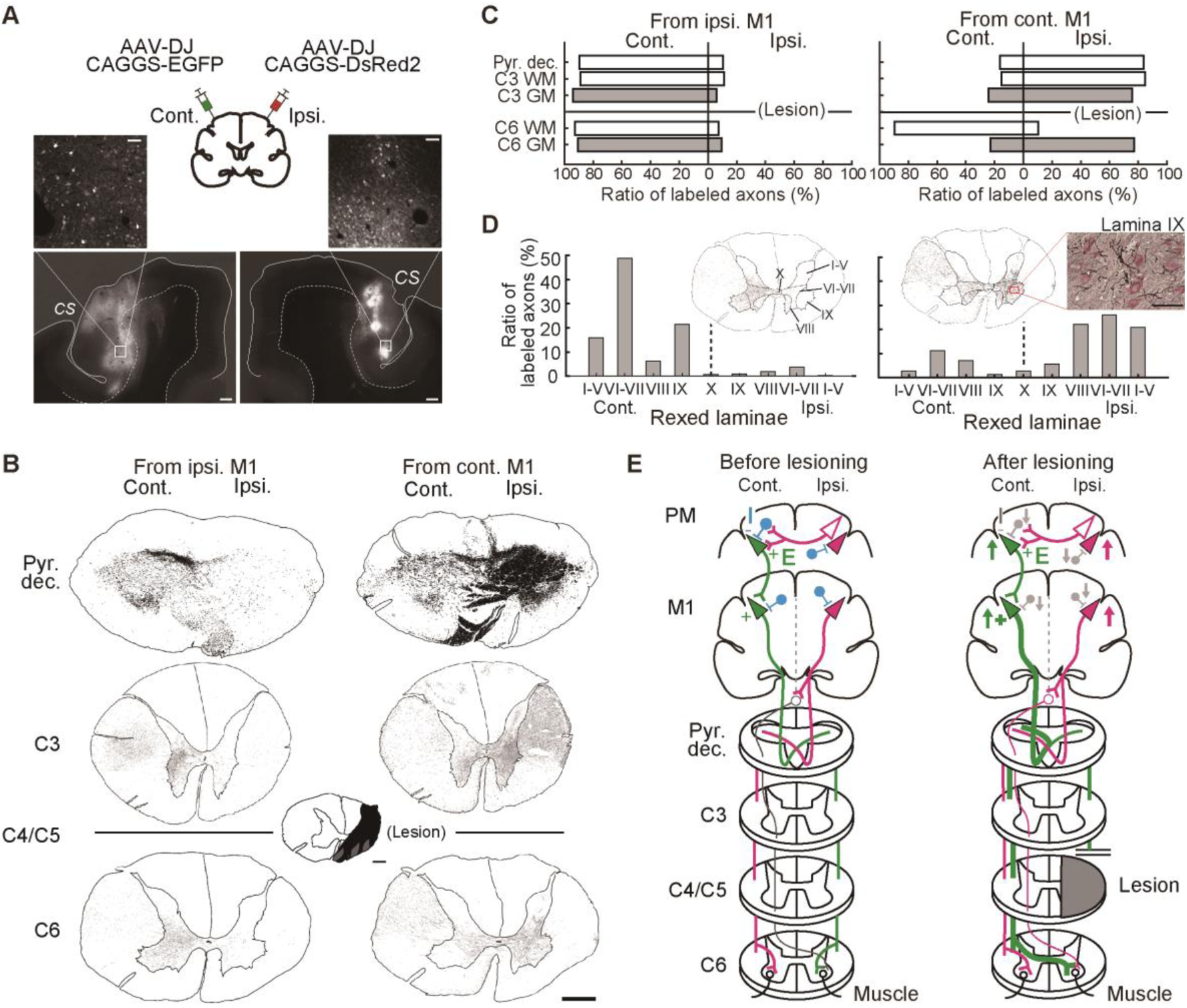
Trajectories of the corticospinal tract after subhemisection in monkey M. (A) Injection sites and labeled cell bodies in the primary motor (M1) area on the contralesional and ipsilesional sides. Scale bars in the lower and higher magnification images are 1,000 and 100 µm, respectively. (B) Distribution of labeled axons at different segments from the pyramidal decussation (Pyr. dec.), C3, and C6. Scale bars: 1,000 µm. The left and right columns show the distribution of labeled axons originating from the ipsilesional and contralesional M1, respectively. (C) The proportion of labeled axons between the contralesional and ipsilesional sides in each segment. At C3 and C6, the white and gray bars indicate the proportion of labeled axons in the white matter (WM) and gray matter (GM), respectively. The proportions of the number of labeled axons between the contralesional and ipsilesional sides was calculated by dividing the number of labeled axons in each side by the all labeled axons in each segment (pyramidal decussation, WM in C3, GM in C3, WM in C6 and GM in C6), expressed as a percentage. (D) Distribution of labeled axons in the Rexed laminae at C6. Regions of interest (ROIs) were classified into nine areas (I–V, VI–VII, VIII, and IX on both sides and X). The proportion of the number of labeled axons in the nine ROIs was calculated by dividing the number of labeled axons in each ROI by all labeled axons in GM at C6. The high magnification image shows that the labeled axons originating from the contralesional M1 terminate at the motoneurons in lamina IX on the ipsilesional side. Scale bar: 100 µm. (E) Schematic diagram of the hypothesis on the switch from interhemispheric inhibition to facilitation. Before lesioning, the commissural cortical neurons excite the target neurons on the contralateral side with direct excitatory connections and inhibit them through inhibitory interneurons (blue). The green and magenta lines represent the corticospinal tract originating from the contralesional and ipsilesional sides, respectively. PM, premotor; M1, primary motor. After lesioning, bilateral motor-related areas are activated and some modulatory inputs reduce the excitability level of the inhibitory interneurons (gray), then the interhemispheric interactions become facilitatory.

## Discussion

Fig. 5E shows a schematic diagram of the hypothesis for the recovery mechanisms of the animals in the present study. The commissural cortical neurons may directly excite the target neurons on the contralateral side and inhibit them through inhibitory interneurons (*19*). After lesioning, some modulatory inputs may reduce the excitability level of the inhibitory interneurons, by which the E/I balance was changed to be more excitable, resulting in switching from interhemispheric inhibition to interhemispheric facilitation. Presumably, such a change in the E/I balance occurs globally and in a nonspecific manner across wide cortical areas at least spanning the PM, M1, and S1. Such global disinhibition might have been effective in inducing muscle twitches in the intact and affected forelimbs from wide cortical areas as shown in Fig. 3. Global disinhibition may open the workspace for the brain to seek an appropriate solution for reorganizing the sensorimotor circuits that were not affected directly by the lesion for functional recovery. This would be one of the basis for diaschisis to promote the recovery. Such a change in interhemispheric interaction has been reported in patients with neurological disorders (*20*) and in the aged populations (*21*), and could be a general compensatory mechanism for decreased motor function. A key question is the origin of the modulatory inputs that change the excitability level of the inhibitory interneurons. The increase in excitability observed in this study cannot be explained by low-level mechanisms such as spasticity, that is, the enhanced stretch reflex often observed after the spinal cord injury. We did not clinically observe spasticity nor increased muscle tone in our monkeys as diagnosed by a specialist neurologist. At this stage, the monkeys became able to perform reach and grasp movements smoothly (Fig. 1 and Movies S1 and S2). Instead, the ECoG responses in the PM/M1/S1 to stimulation of the contralateral side were enhanced (Fig. 3), which supported the cortical origin of excitability. Recent studies suggest that vasoactive intestinal peptide (VIP) interneurons mediate disinhibition in the neocortical circuits and their activation promotes recovery after stroke (*22*). Considering its global and nonspecific properties, it is likely to be related to subcortical divergent projection systems such as cholinergic (*23*), monoaminergic (*24*) or serotonergic (*25*) neurons, all of which modulate the GABAergic interneurons in the cortex including the VIP interneurons. The role of glial cells in this process might also be claimed and needs to be tested (*26*).

Previous studies of macaques with subhemisection showed much slower and poorer recovery compared to our animals (*12, 13*), even though it is better than observed in rodents (*27*). Another key factor supporting the drastic recovery in the animals of the present study would be the massive plasticity of the CST originating from the contralesional M1 (Figs. 5 and S25), to an extent that has never been reported. Considering the experimental conditions between the preceding studies and current study, possible factors that induced such massive plasticity would be the intensive training and extensive electrical stimulations of the bilateral PM, M1 and S1 applied through the ECoG electrodes to test for muscle twitches (3mA × 3 pulses × 100 trials × 36 or 56 Ch /week). The effect of electrical stimulation of the motor cortex on sprouting of the corticospinal tract and promoting post-injury recovery has been demonstrated in a rodent model (*28*). Clarifying the mechanism of global disinhibition and massive plasticity of corticospinal axons may give us insights for developing therapeutic strategies to promote functional recovery after severe neuronal injury.

In the present study, we performed subhemisection of the spinal cord, but later found that some tissue seemed to have remained as “islands” in the lesioned area, which contained large number of glial cells and where Klüver-Barrera staining was faint. We believe the lesion was complete, as shown in Fig. S3, but there is a possibility that axon bundles regenerated during recovery course. Similar observations were reported by Cavada and colleagues in which macaques with complete thoracic spinal cord injury showed recovery in hindlimb function and corticospinal conduction to hindlimb muscles at 11–13.5 months after injury (*29*). The possibility of such extensive regeneration should be examined in future studies.

## Acknowledgements

We thank Jiro Yamashita, Masashi Nakamura, Yuta Shinto and Erika Omae for technical assistance.

## Funding

This work was supported by a Grant-in-Aid for Scientific Research on Innovative Areas “Hyper-adaptability” to TI (Project no. 19H05723), a Grant-in-Aid for Scientific Research from the Ministry of Education, Culture, Sports, Science, and Technology (MEXT) to TI (KAKENHI (A) no. 19H01011 and (S) no. 22H04992), Japan Agency for Medical Research and Development (JP18dm0307005 to NS and TI), a Grant-in-Aid for Scientific Research from MEXT to RY (KAKENHI (B) no. 21H02798) and a Grant-in-Aid for Scientific Research from MEXT to HO (KAKENHI (A) no. 20H00573).

## Author contributions

Conceptualization: RY, SU, TI

Data curation: RY, SU, TK

Formal analysis: RY, SU, TK, MM, ZC, TT

Funding acquisition: RY, TI

Investigation: RY, SU, TK, MM, KI

Methodology: RY, SU, KI, KK, JT, HO, TI

Project administration: RY, TI

Resources: RY, KK, JT, HO, TI

Software: RY, ZC

Supervision: ZC, TT, TI

Validation: TI

Visualization: RY, SU, KI

Writing - original draft: RY, SU, TI

Writing – review & editing: RY, SU, TI

## Competing interests

The authors declare that they have no competing interests.

## Data and materials availability

The data and codes supporting the findings of this study are available from the corresponding authors upon reasonable request.

## Supplementary Materials

### Materials and Methods

#### Subjects

Two macaque monkeys (*Macaca fuscata*, monkeys M and H, both male, bodyweight 6.5 and 5.2 kg, respectively) were used in the present study. The experimental protocols followed the guidelines set forth by the Ministry of Education, Culture, Sports, Science and Technology of Japan, and were approved by the Institutional Animal Care Committee and Kyoto University, Japan.

#### Task training

To assess finger dexterity, the monkeys were trained to perform a reach-and-grasp task (Fig. 1A). The subjects were seated in a monkey chair with their heads fixed and were required to put their right hand on a board positioned at the height of their abdomen. A cube of sweet potato (6 × 6 × 6 mm) was presented between a 9-mm-wide vertical slit after the monkeys pressed a button with their right hand for at least for 2 s, and the subjects could the retrieve the piece of food. One hundred trials were recorded on each experimental day. A successful precision grip was identified as the successful removal of the sweet potato cube using the pads of the index finger and thumb. A digital video camera (33 frames/s) was used to record the reach-and-grasp sequence from the lateral view. A successful trial was defined as any trial that resulted in the removal of the food from the slit without dropping it. Forelimb movements were recorded using an optical motion capture system that used 12 infrared cameras (Eagle-4 Digital RealTime System, Motion Analysis; Motion Analysis, Rohnert Park, CA, USA). The spatial positions of the reflective markers (6-mm-diameter spheroids) were sampled at 200 Hz. Nineteen markers were attached to the surface of the forelimb using double-sided tape at the following positions: center of the right chest (marker 1 [m1]), proximal and distal parts of the right upper arm (m2 and m3), proximal and distal parts of the right fore arm (m4 and m5), back of the right hand (m6, m7, and m8), first and second joints of the thumb (m9 and m10), first, second, and third joints of the index finger (m11, m12, and m13), first, second, and third joints of the index finger (m14, m15, and m16), and first, second and third joints of the fifth finger (m17, m18, and m19). Reaching onset was measured as the time when the right hand was lifted from the button; grasping onset was measured as the time when the tip of the index finger (m14) was inserted in the slit.

#### Surgical procedures

All surgeries described below were performed using sterilization under general anesthesia, starting with a combination of ketamine (10 mg/kg, intramuscular injection [i.m.]) and xylazine (1 mg/kg, i.m.) and succeeded by intubation and isoflurane (1–1.5%) inhalation to maintain stable, deep anesthesia throughout surgery. Heart rate, peripheral capillary oxygen saturation and end-expiratory carbon dioxide pressure were monitored during surgery. Ringer’s solution was administered continuously through an intravenous (i.v.) drip. Dexamethasone (0.825 mg/ kg body weight) and ampicillin (40 mg/kg) were administered after anesthesia.

##### Implantation of the head post holder, EMG wires, and ECoG electrode array

Titanium head post holders were implanted on the skull in aseptic conditions. EMG activitiy of the right forelimb muscles was recorded through chronically implanted pairs of multi-stranded stainless steel wires (Cooner Wire, Chatsworth, CA, USA), which were tunneled subcutaneously to their target muscles. Circular connectors (MCP-12, Omnetics, Minneapolis, MN, USA) were anchored to the skull. EMGs were recorded from selected forearm muscles. In monkey M, EMG was recorded from brachioradialis (BR), triceps brachii (TRI), biceps brachii (BB), extensor digitorum communis (EDC), extensor digitorum 2 and 3 (ED23), extensor carpi ulnaris (ECU), flexor digitorum superficialis (FDS), flexor carpi radialis (FCR), flexor carpi ulnaris (FCU), palmaris longus (PL), adductor pollicis (ADP) and first dorsal interosseous (FDI). In monkey H, EMG was recorded from the middle deltoid (DM), BR, TRI, BB, EDC, ED23, ECU, FDS, FCR, FCU, PL, ADP, and FDI. EMG signals were amplified, filtered (5–1,000 Hz), and rectified. All EMG signals were sampled at 2,000 Hz.

Cortical oscillatory activity was recorded from the sensorimotor cortex, including the premotor cortex (PM), primary motor cortex (M1), and somatosensory cortex (S1). The skull was exposed over the bilateral frontal cortices. Craniotomies were located around the central sulcus, and the cortex around the central sulcus was exposed bilaterally. Two platinum ECoG arrays, each comprised of 18-channel (6 × 3 grid) electrodes in monkey M and each comprised of 28-channel (7 × 4 grid) electrodes in monkey H, were placed on the digit, hand, and arm areas of the M1, PM and S1 on both hemispheres (Fig. 1B). The electrodes had a diameter of 1 mm and an inter-electrode distance of 3 mm center-to-center. Small plastic screws were attached to the skull as anchors. To estimate the motor areas covered by the ECoG electrodes, a microstimulation current through a stimulator (Nihon Kohden, Tokyo, Japan) was delivered to each ECoG electrode on the contralesional hemisphere, and movements were examined. Each electrode was labeled based on the body parts of which movements could be evoked. To verify the position of the ECoG arrays, the brain was removed at the end of all experiments after the transcardial perfusion of 4% paraformaldehyde, photographed, and the electrode locations were reconstructed. The electrode locations were further estimated by comparing magnetic resonance images and the physical dimension of the arrays.

ECoG recordings that were used in the analysis of cortical activation and corticocortical interactions were made on different days before and after spinal cord lesioning. In monkey M, data were recorded on Days −10, −9, −8, −2, −1, 32, 33, 36, 37, 39, 40, 43, 45, 49, 52, 54, 57, 58, 60, 61, 64, 71, 72, 74, 78, 79, 81, 82, 84, 87, 89, 92, 93, 95, 96, 100, 103, 106, 107, 110, 114, 117, 120, 122, 130, 133, 137, 138, 140, 142, 145, 148, 151, 152, and 154. In monkey H, data were recorded on Days −5, −4, −2, 23, 24, 25, 28, 31, 32, 35, 37, 38, 39, 43, 44, 45, 46, 50, 51, 52, 53, 56, 58, 59, 60, 65, 67, 70, 74, 77, 81, 84, 85, 87, 88, 91, 94, 98, 100, 102, 107, 108, 112, 114, 116, 119, 121, 122, 123, 128, 129, 130, 133, 134, 135, 137, 140, 141, 142, 143, 147, 149 and 150. Signals were recorded with a Plexon MAP system (Plexon, Dallas, TX, USA) at a sampling rate of 2,000 Hz. ECoG signals were extracted using multichannel amplifiers with 0.3 Hz high-pass and 7,500 Hz low-pass analog filters.

##### Spinal cord injury (SCI)

The subhemisection was made in both monkeys after pre-lesion data were obtained. Under the above-mentioned anesthesia, the border between the C4 and C5 segments (C4/C5) was exposed by laminectomy of the C3 and C4 vertebrae, and a transverse opening was made in the dural membrane. After identification of the dorsal roots at the C4 and C5 levels, the border between the C4 and C5 segment was lesioned with a surgical blade and a micro-tweezer. In this SCI model, the lateral funiculus and a part of the ventral and dorsal funiculi were injured. The skin and back muscles were sutured with nylon or silk.

#### Data analysis

##### Behavior

The number of successful trials according to the type of grasping in each experimental day was recorded. The three following types of grasping were defined. If the monkeys could take the food from the slit using the pads of the index finger and thumb without being aided by the slit, this movement was judged as a precision grip. If the monkeys could take the food from the slit by being aided by the slit, this was judged as a pinch along the slit. In the present study, the monkeys were trained using the task without the slit after subhemisection because the monkey could not move the fingers independently just after subhemisection. If the monkeys could take the food from the devices without a slit, the movement was judged as a gross power grip.

##### Preprocessing for brain activity

The 60-Hz line noise was removed from the raw ECoG data using the MATLAB FieldTrip toolbox (*30*). The data were then downsampled four times, resulting in a sampling rate of 500 Hz. Channels and trials with abnormal spectra were rejected using an automated algorithm from the EEGLAB library (*31*), which has been suggested as the most effective method for artifact rejection (*32*). The ECoG signals from each channel were then aligned using the timing of reaching onset (Time 0), where grasping onset varied but typically occurred at 0.3–2 s (monkey M: 0.3–2 s; monkey H: 0.3–0.6 s) after reaching onset. Further analyses of cortical activation and corticocortical interactions were performed using the MATLAB FieldTrip toolbox.

##### Dynamics of cortical activation

The dynamics of cortical activation in each channel were quantified by the time-frequency representation (TFR) generated by Morlet wavelet transformation, at 113 different center frequencies (8–120 Hz) with the half-length of the Morlet analyzing wavelet set at the coarsest scale of five samples (Fig. 2B). The α band was defined as 8–12 Hz, β band as 13–30 Hz, γ band as 31–70 Hz and high-γ band as 71–120 Hz. To quantify event-related activity (ERA) during reaching-and-grasping, we further divided each TFR value by the baseline value (the mean TFR value at the corresponding frequency during the resting period from −1.5 to −1 s).

##### Dynamics of corticocortical interactions

The dynamics of corticocortical interactions were quantified by spectral Granger causality (GC), which can represent phase differences between signals from two cortical areas to provide their asymmetric causal dependence (*33, 34*). Between signals from two channels, in-trial spectral GCs were calculated for each experimental day, where each GC represents a unidirectional connectivity from one channel to another and across frequencies between 8 and 120 Hz (113 frequency bins). Event-related causality (ERC) was further quantified by dividing each GC value by the baseline value (the mean GC value at the corresponding frequency during the resting period from −1.5 to −1 s).

Three preparation steps were performed for spectral GC calculation.

1. Preprocessing: detrending, temporal normalization, and ensemble normalization were performed to achieve local stationarity of the data (*35*). Detrending, which is the subtraction of the best-fitting line from each time series, removes the linear drift in the data. Temporal normalization, which is the subtraction of the mean of each time series and division by the standard deviation, ensures that all variables have equal weights across the trial. These processes were performed on each trial for each channel. Ensemble normalization, which is the pointwise subtraction of the ensemble mean and division by the ensemble standard deviation, dramatically improves the local stationarity of the data (*35, 36*).
2. Window length selection: the length and step size of the sliding-window for segmentation were set as 150 and 20 ms, respectively.
3. Model order selection: model order, which is related to the length of the signal in the past that is relevant to the current observation, was determined by the Akaike information criterion (AIC) (*37*). In both subjects, a model order of 10 samples (equivalent to 10 × 4 = 40 ms of history) resulted in a minimal AIC and was selected. The selected model order also passed the Kwiatkowski–Phillips– Schmidt–Shin (KPSS) test (*38*), thus maintaining local stationarity. Furthermore, the vector autoregression model was validated by a consistency test (*35*).

#### Electrical stimulation through ECoG electrodes

ECoG electrodes delivered a microstimulation current through a stimulator (Nihon Kohden, Tokyo, Japan) to evoke muscle twitches in the forearm. During a stimulation sequence, 3 pulses with an interval of 50 ms were repeated 100 times (intensity of a single monophasic current pulse: 3 mA, pulse duration: 0.5 ms; inter-trial interval: 2–4 s). The monkeys needed to sit in the monkey chair and maintain a resting state without any movements. If the monkeys moved, those trials were excluded from the analysis. Cortical stimulation was delivered at 18 channels in both hemispheres every week after lesioning. Muscle twitches in the forearm and evoked potential on the opposite hemisphere to the stimulation side were recorded. For the muscle twitches in each body part induced by electrical stimulation, the magnitude of the twitches was judged by the experimenter and classified into four ranks (3 = muscle twitches with joint movements, 2 = visible muscle twitches, 1 = invisible muscle twitches, 0 = no response). Electrical stimulation was applied from channels 1 to 18 in monkey M and channels 1, 2, 3, 5, 6, 7, 9, 10, 11, 13, 14, 15, 17, 18, 19, 21, 22, and 23 in monkey H (Fig. S1).

#### Conditioning-test

To investigate the effect of inter hemispheric interactions on muscle activity, a conditioning-test approach was utilized. A conditioning stimulus was used to activate the ipsilesional PM without the activation of muscle activity on the affected side. After conditioning stimulation, a test stimulus was used with several time intervals to induce the muscle activity of the affected side. The intensity of the conditioning and test stimuli was sub-threshold (0.9 T) and supra-threshold (1.1 T), respectively. The inter-stimulus intervals (ISI) between the conditioning and test stimuli were 0, 5, 10, 15, 20, 25, 30, 40, and 50 ms. To compare the effect of the conditioning stimulus, muscle activity was recorded with test stimulation only.

#### Viral vector preparation

The adeno-associated virus (AAV) vectors were prepared as described previously (*39*). Briefly, the packaging (pAAV-DJ and pHelper) and transfer plasmids (pAAV-CAGGS-EGFP and pAAV-CAGGS-DsRed2) were transfected into HEK293T cells. After incubation, the harvested cells were lysed and purified by serial ultracentrifugation with cesium chloride. The purified particles were dialyzed with 0.001% Pluronic-F68 Solution (Sigma-Aldrich, St. Louis, MO, USA) in phosphate-buffered saline (PBS), and then concentrated through ultrafiltration. The copy number of the viral genome (vg) was determined using a TaqMan Universal Master Mix II (Applied Biosystems, Foster City, CA, USA).

#### Viral vector injections

After the behavioral observation period, two different AAV vectors were injected into the M1 area of each hemisphere. Under isoflurane anesthesia, AAV DJ-CAGGS-EGFP or AAV DJ-CAGGS-DsRed was injected using a glass pipette (outside diameter at tip, 60–110 µm) attached to a Hamilton syringe (10 μL, Model 701, Cemented NDL, 26sG, 2 in, point style 2) at multiple locations. In monkey M, AAV DJ-CAGGS-EGFP (titer, 1.5 × 10^13^ vg/ml) was injected at 24 locations in the contralesional M1 as follows: at four depths (1.5, 3, 4.5 and 6 mm from the cortical surface) in four penetration tracks at approximately 1mm anterior to the CS, and at one depth (1.5 mm) in four tracks approximately 1.5 and 3 mm anterior to the preceding injections. Medio-laterally, each injection was separated by 1.5 mm. The actual injection of the vector was successively conducted after a 5-min interval to allow for stabilization of the tissue after needle insertion. Subsequently, 0.5 µl vector was injected at a rate of 0.1 µl/min. Following the initial injection, another 5-min interval ensured that the viral vector was absorbed, and then, the needle was moved slowly. In addition, AAV DJ-CAGGS-DsRed (Titer, 1.6 × 10^13^ vg/ml) was injected into the ipsilesional M1 at the same coordinates as for the contralesional M1. To validate the difference of the virus vectors, the vectors were switched between monkeys H and M: AAV DJ-CAGGS-DsRed (Titer, 2.4 × 10^13^ vg/ml) was injected into the contralesional side and AAV-CAGGS-EGFP (titer, 2.4×10^13^ vg/ml) was injected into the ipsilesional side. Each virus vector was injected at 21 locations (4 depths × 4 tracks, 1 depth × 5 tracks).

#### Immunohistochemistry

At 7 weeks after the last injection, the monkeys were anesthetized deeply and perfused transcardially with 4 % paraformaldehyde in 0.1 M phosphate buffer. The brain and spinal cord were removed and immersed in sucrose following post-fixation. Coronal sections of the brain including the injection area and transverse sections from the medulla to the thoracic segment were prepared (40 μm thick). Some sections including the lesion area were processed for Klüver-Barrera staining.

Additional sections of the medulla to the thoracic level were stained by immunohistochemistry to visualize the GFP or RFP labeled axons. Free-floating sections were washed three times in 0.05M PBS for 10 min each and incubated with 0.6 % hydrogen peroxide in methanol for 30 min. Following the wash with PBS, the sections were blocked in 5 % skim milk in PBS with 0.3 % Triton X-100 (PBST) for 60 min. The sections were then incubated in rabbit anti-GFP antibody (1:4,000; A11122, Invitrogen, Carlsbad, CA, USA) in PBST for 60 min at room temperature following overnight at 4 ℃. After washing four times with PBST for 5 min, the sections were incubated with a biotinylated goat anti-rabbit IgG antibody (1:200; BA-1000, Vector Laboratories, Burlingame, CA, USA) for 2 h. Following washing with PBST, the sections were treated with PBST containing avidin-biotin-peroxidase complex (ABC Elite, 1:100; PK-6100, Vector Laboratories) for 60 min. The sections were washed three times for 5 min in PBS and TBS in turns and treated with TBS containing 0.01 % diaminobenzidine (DAB; 040-27001, FUJIFILM Wako Pure Chemical Corporation, Tokyo, Japan), 1.0 % nickel-ammonium sulfate (FUJIFILM Wako Pure Chemical Corporation, 146-01012), and 0.0003 % hydrogen peroxide. The sections were washed three times in TBS, three times in PBS for 5 min each.

To examine RFP expression, free-floating sections were washed three times in PBST for 10 min and blocked in PBST containing 10 % normal goat serum (S-1000, Vector Laboratories). A rabbit anti-RFP antibody (1:2,000; 600-401-379, Rockland, Immunochemicals, Inc., Boyertown, PA, USA) was applied for 60 min at room temperature and then overnight at 4 ℃. Following washing four times with PBST for 5 min, the sections were incubated with a secondary antibody (1:200; BA-1000, Vector Laboratories) for 2 h. After washing four times in PBST for 5 min, the sections were treated with ABC Elite (1:200; PK-6100, Vector Laboratories). Washing and reaction with DAB solution were conducted as same in the previous protocol for anti-GFP.

After being mounted on gelatin-coated glasses, some sections were counterstained with Neutral Red. Digital images were acquired using a BZ-X710 Keyence microscope. The labeled axons were drawn using a camera lucida attached to a BZ51 Olympus microscope with a 20× objective lens.

For immunofluorescence, free-floating sections were washed three times in PBS with 0–0.6 % Triton X-100 (PBST) for 10min and blocked with PBST containing 10% normal goat serum (NGS) for 60 min. A rinse step with PBST containing 2% NGS for 10 min was performed only for anti-GFP reactions. The sections were incubated for 60 min at room temperature and overnight at 4 ℃ with a rabbit anti-GFP antibody (1:2,000; A11122, Invitrogen), or rabbit polyclonal anti-RFP antibody (1:1000; 600-401-379, Rockland Immunochemicals, Inc.). After washing three or four times in PBST with 2 % NGS for 10 min, anti-rabbit IgG AF647 (1:200; A21244, Invitrogen) in PBST with 2 % NGS was applied and incubated for 3 h in the dark. The sections were washed four times with PBS for 5 min and mounted on gelatin-coated glass slides. Images were acquired using a BZ-X710 Keyence microscope.

#### Quantification of labeled axons

For the pyramidal decussation, the regions where corticospinal axons descend to the caudal segments were separated for quantification. The medial part was not included because many axons projecting to each side are mixed in that region. For the spinal cord, the white matter and gray matter were separated. The sum of the lateral funiculus and ventral funiculus was shown as white matter in this study. To define the borders of the laminae on gray matter, we referred to the cytoarchitecture of Rexed (*40*). The number of traced axons in the immunostaining sections was counted on each part of the sections. When a fiber crossed the border between the regions of both sides, the axon was counted in both regions. The sum of the data on both sides was 100% and the ratio of the labeled axons on the ipsilesional side versus the contralesional side was calculated.

---

**Fig. S1.**
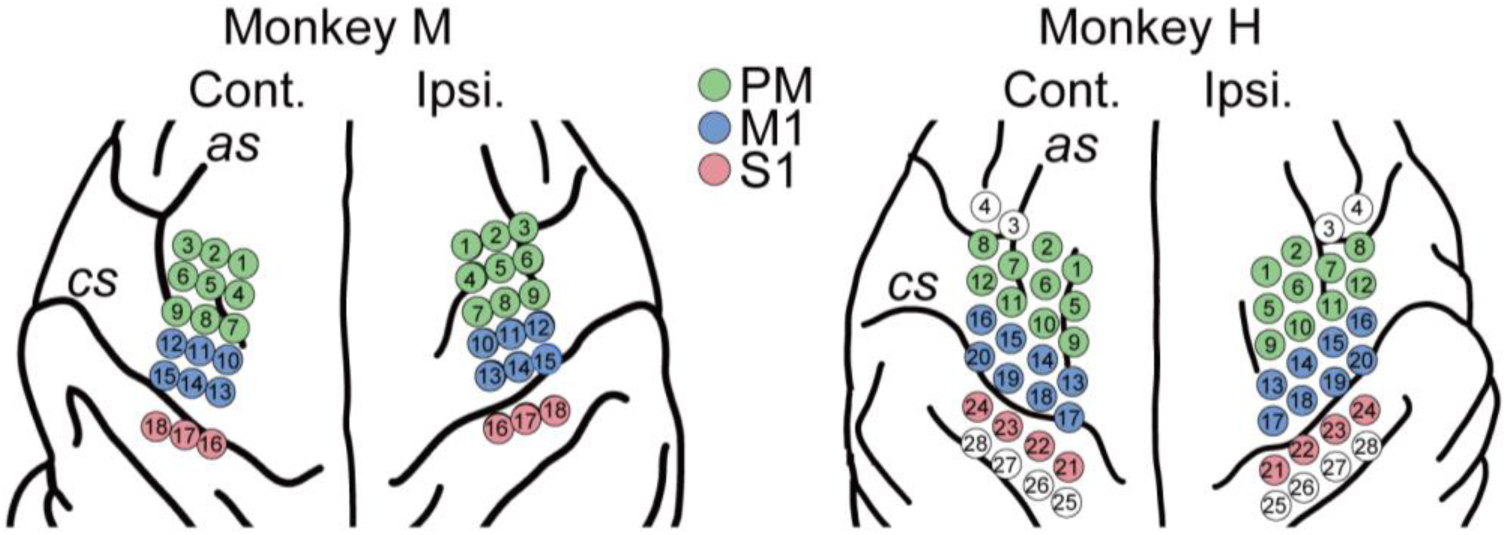
Distribution of electrocorticography channels across the bilateral sensorimotor cortical areas in monkeys M (left) and H (right). as, arcuate sulcus; Cont., contralesional; cs, central sulcus; Ipsi., ipsilesional. In monkey M, channels (Ch) 1–9 were in the premotor (PM) area, Ch10–15 were in the primary motor (M1) area, and Ch16–18 were in the primary somatosensory (S1) area on both sides. In monkey H, Ch1, 2, and 5–12 were in the PM, Ch13–20 were in the M1, and Ch21–24 were in the S1.

**Fig. S2.**
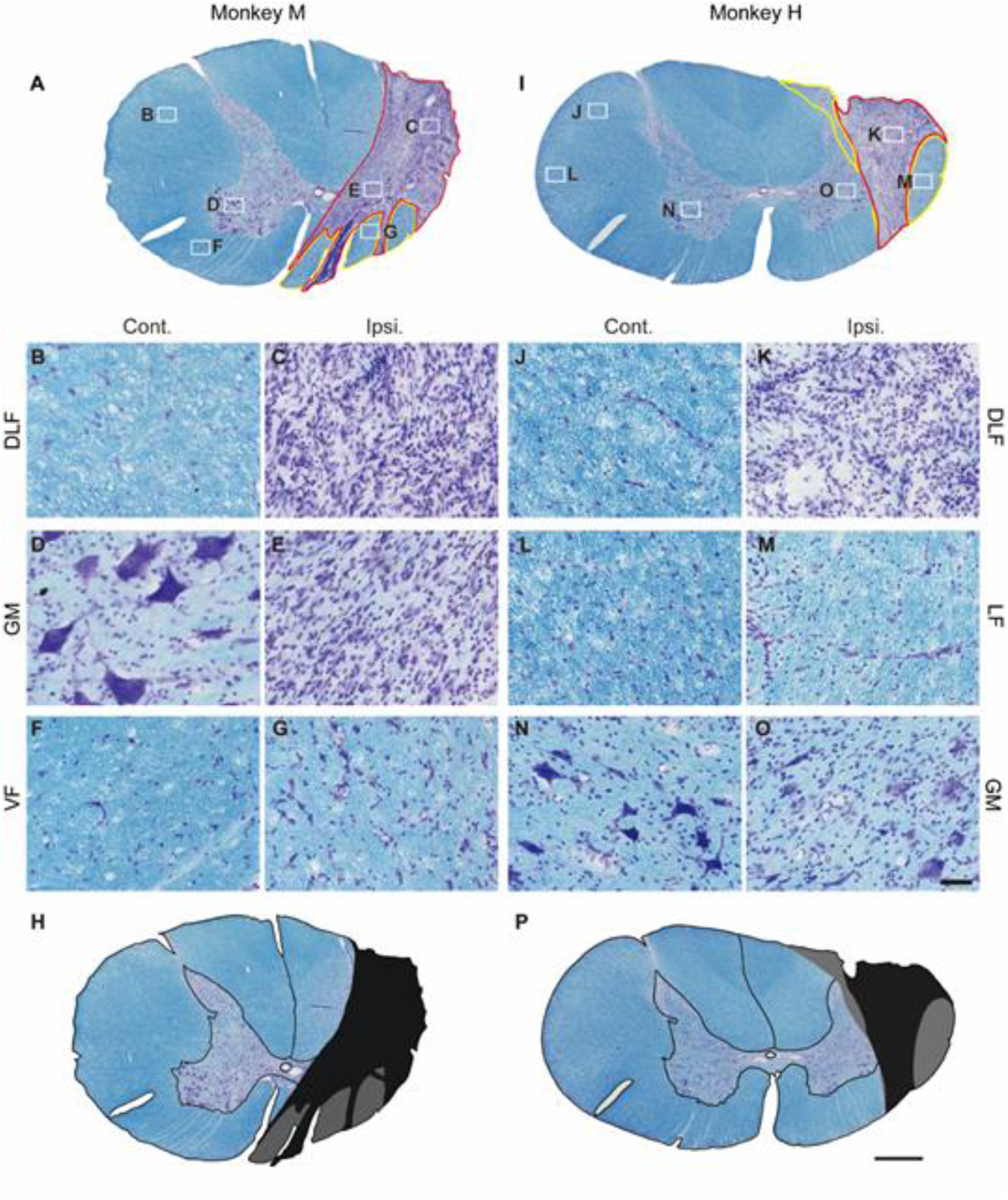
Photomicrographs of spinal cord tissue around the lesioned area with Klüver-Barrera staining. (A) The whole spinal cord at the C4/C5 segment of monkey M stained with Klüver-Barrera. (B–G) Photomicrographs of the areas indicated in (A) with corresponding letters. (H) On the basis of the observations, the lesion extent in monkey M is indicated by the black area. (I–O) The whole spinal cord at the C4/C5 segment was stained with Klüver-Barrera and photomicrographs of the area indicated in (I) with corresponding letters of monkey H. (P) The lesion extent in monkey H is indicated by the black area. Note that the stained images of the dorsolateral funiculus (DLF) where the corticospinal tract descending area is considerably different between the contralesional and ipsilesional sides ([B] and [C] of monkey M, [J] and [K] of monkey H), which means these areas were completely lesioned. In the gray matter (GM) of monkey M, the motoneurons, which are stained clearly on the contralesional side as shown in (D), were not detected on the ipsilesional side as shown in (E), which we also judged as the lesioned area as shown by the black area in (H). Furthermore, the areas indicated with a gray hatch in (H) and (P) contain intact axons; however, as shown in (G) of the ventral funiculus (VF) and (M) of the lateral funiculus (LF), these areas include a number of glial cells and the Klüver-Barrera staining did not look normal. Therefore, we judged that the gray areas are not completely healthy. Since during surgery, as shown in Fig. S3, we confirmed complete transection, there is a possibility that these areas might include reconnected axons. Calibration bars in (A), (H), (I), and (P) indicate 1,000 μm, in (B–G) and (J–O) they indicate 100 µm.

**Fig. S3.**
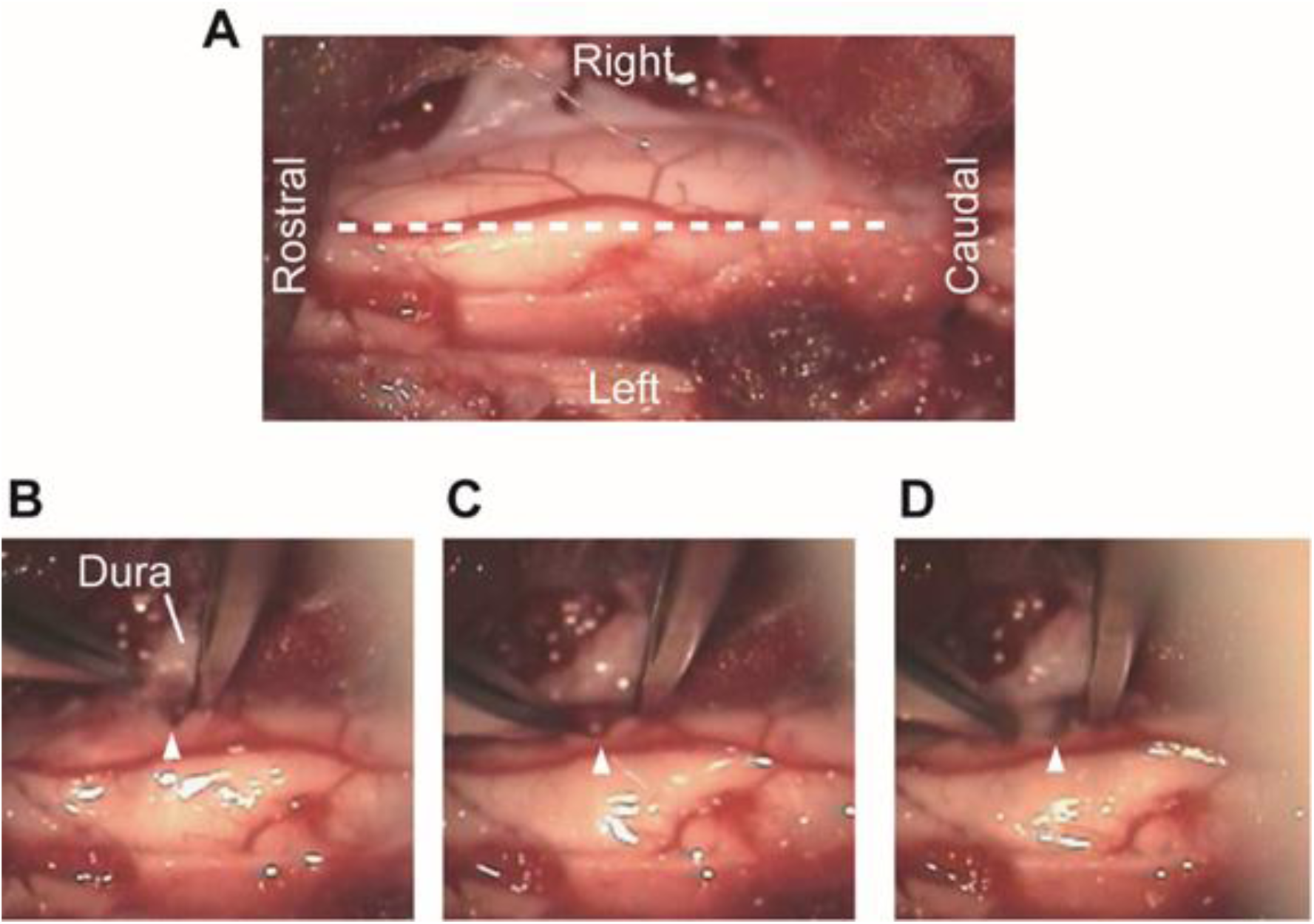
Images from videos taken during surgery for spinal cord subhemisection in monkey H. (A) Low magnification view. (B–D) Higher magnification views with time sequence from (B) to (D). The white triangle indicates the lesion site, which shows that the spinal cord tissue was completely separated at the lesion site.

**Fig. S4.**
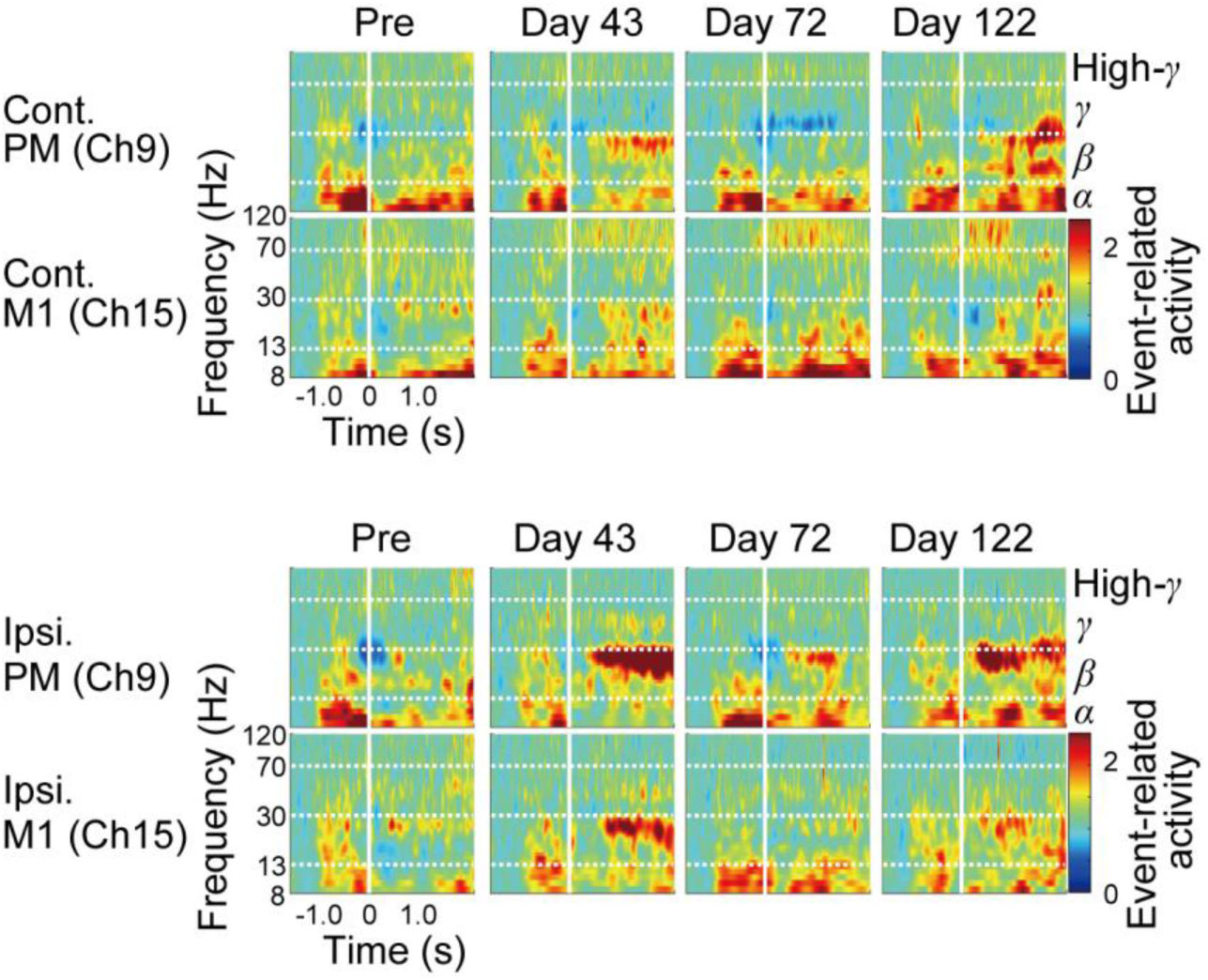
Examples of time-frequency plots of peri-movement activity recorded from the electrocorticography electrodes in the contralesional premotor (PM, channel [Ch] 9) and primary motor (M1, Ch15) areas, and ipsilesional PM (Ch9) and M1 (Ch15) in monkey M before (Pre) and after (Days 43, 72, and 122) lesioning. Time zero on the horizontal axis indicates movement onset.

**Fig. S5.**
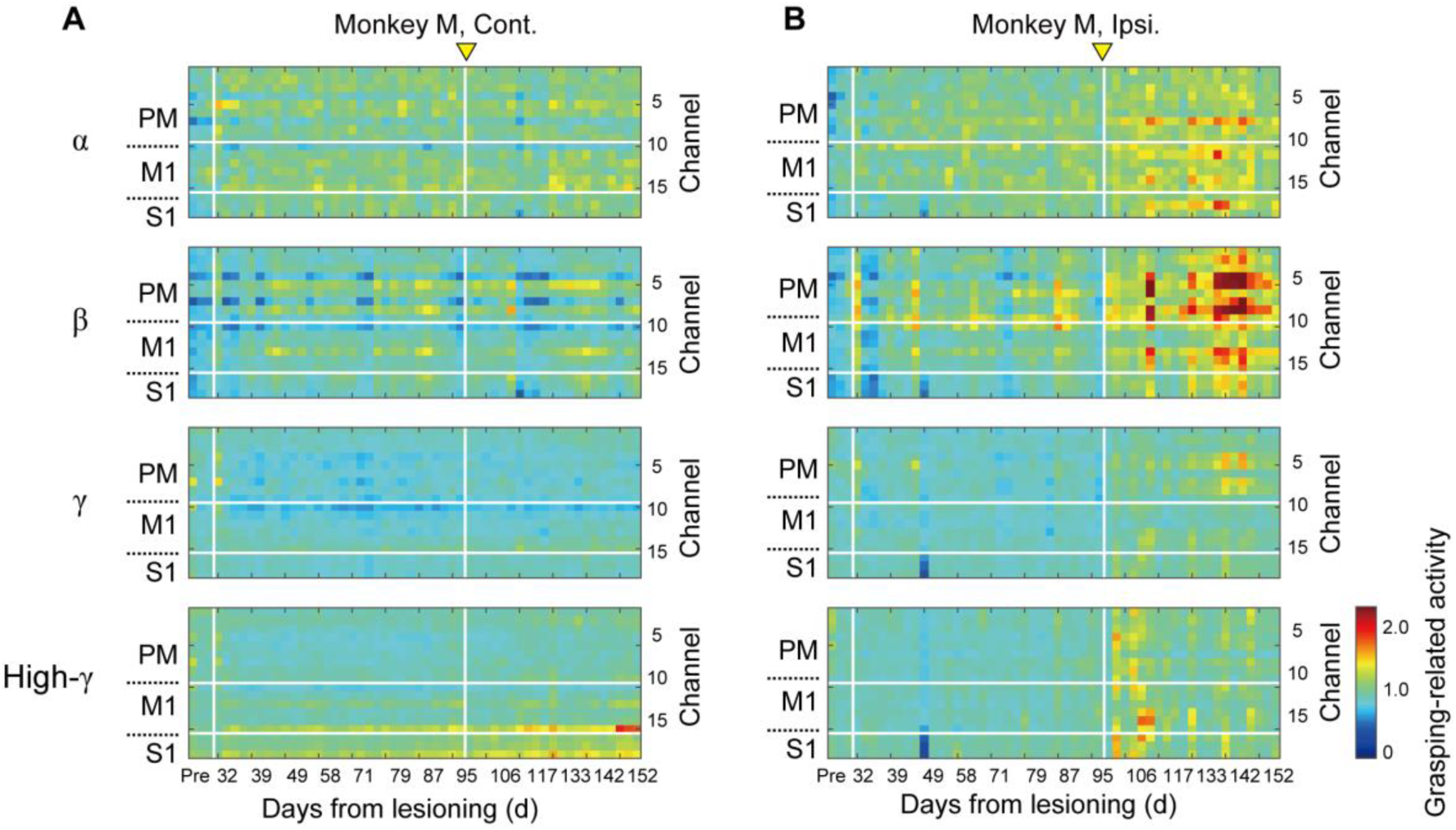
Longitudinal analysis of the magnitude of peri-movement electrocorticography activity at different frequency ranges (α, β, γ, and high-γ) in individual channels in the contralesional (A) and ipsilesional (B) premotor (PM), primary motor (M1), and primary somatosensory (S1) areas before (Pre) and after (Days 32–152) lesioning in monkey M. The vertical white line on Day 95 indicates the timing when recovery was saturated (see Fig. 1D–F). Note that the α, β, and high-γ band activity of the ipsilesional PM and M1 was temporarily enhanced around Day 95.

**Fig. S6.**
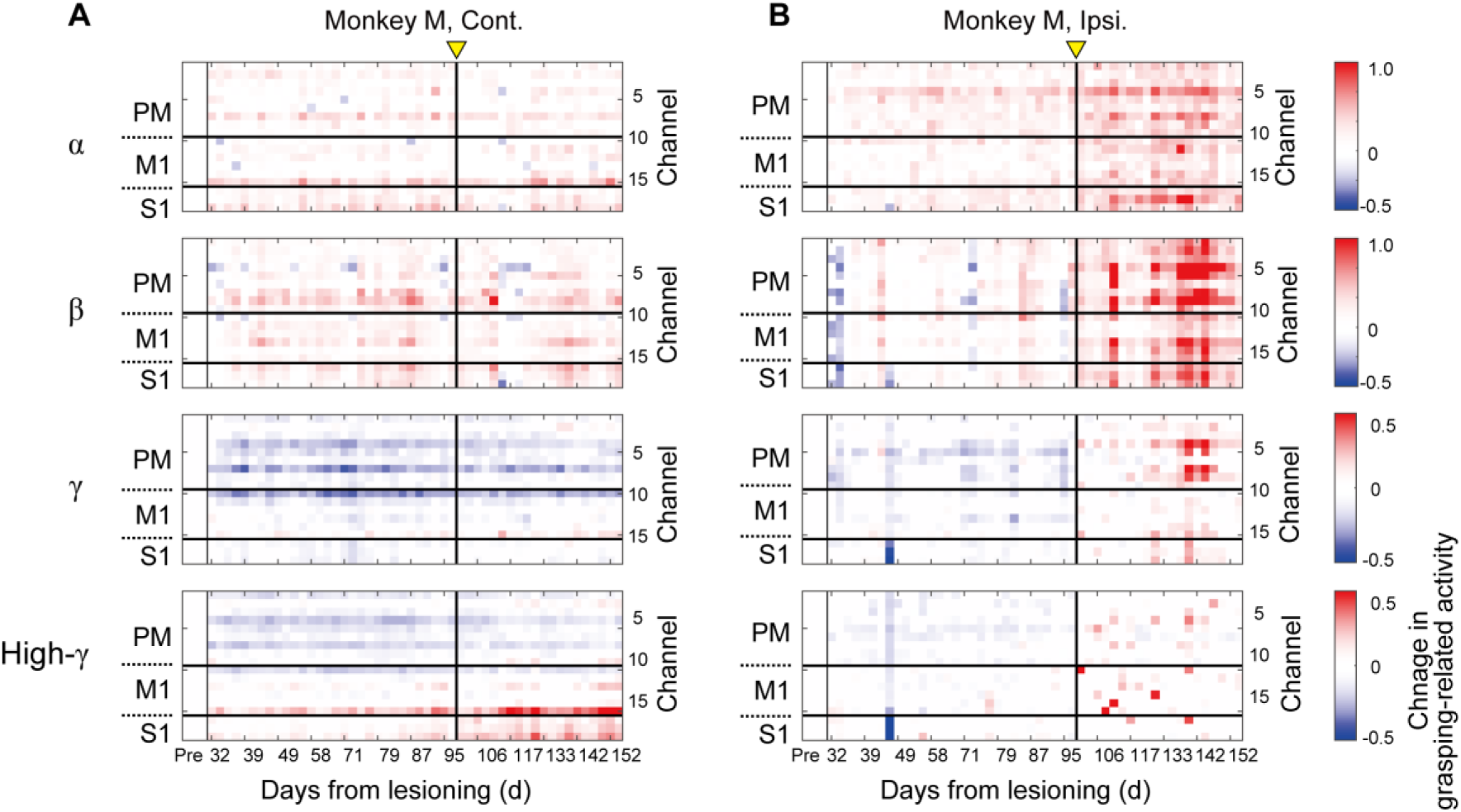
Longitudinal change in the difference of peri-movement electrocorticography activity after the spinal cord injury shown in Fig. S5 from the prelesion data in monkey M. M1, primary motor; PM, premotor; S1, primary somatosensory.

**Fig. S7.**
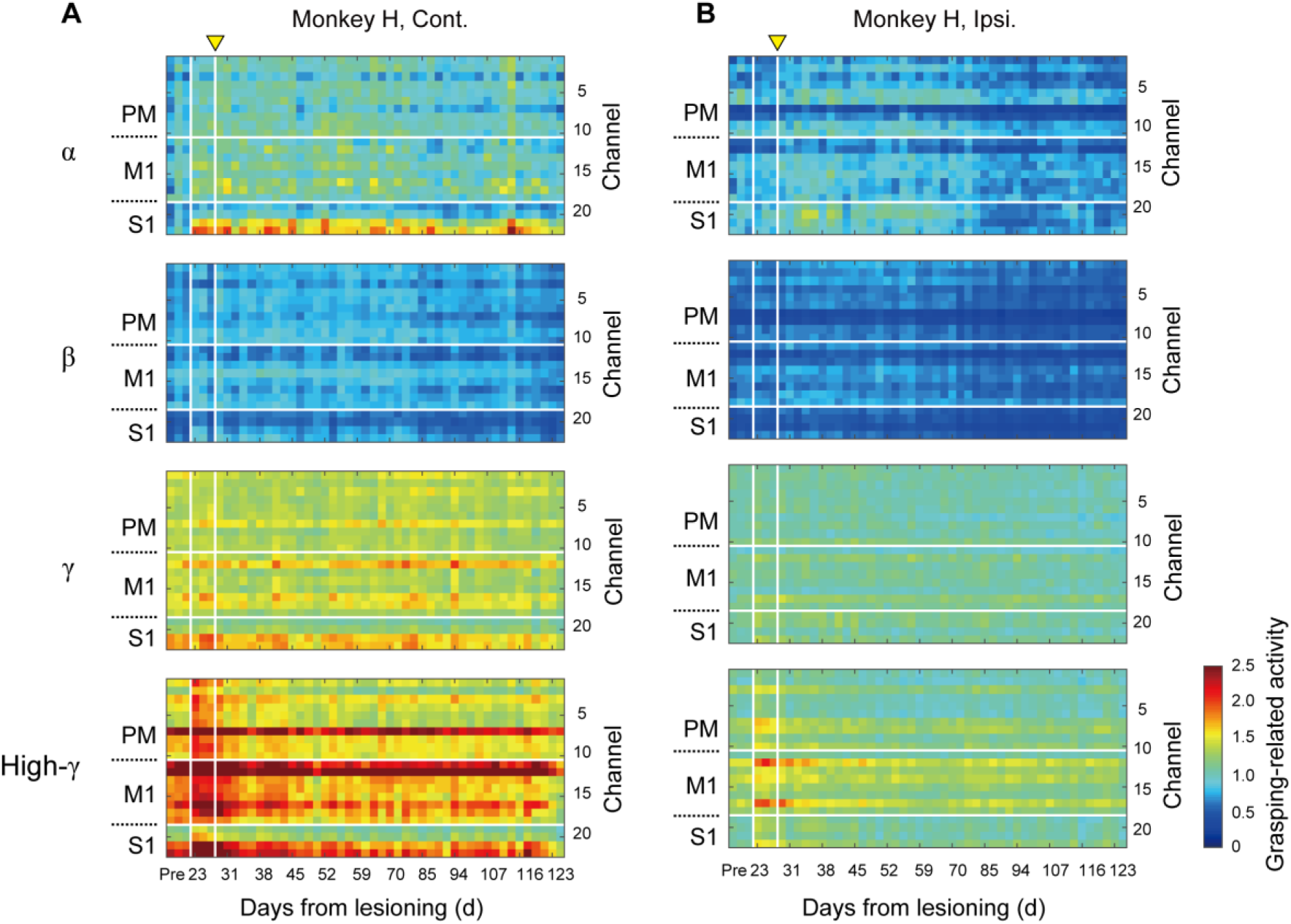
Longitudinal analysis of the magnitude of peri-movement electrocorticography activity at different frequency ranges (α, β, γ, and high-γ) in individual channels in the contralesional (left) and ipsilesional (right) premotor (PM), primary motor (M1), and primary somatosensory (S1) areas before (Pre) and after (Days 23–123) lesioning in monkey H. The vertical white line on Day 27 indicates the timing when recovery was saturated (see Fig. 1D–F).

**Fig. S8.**
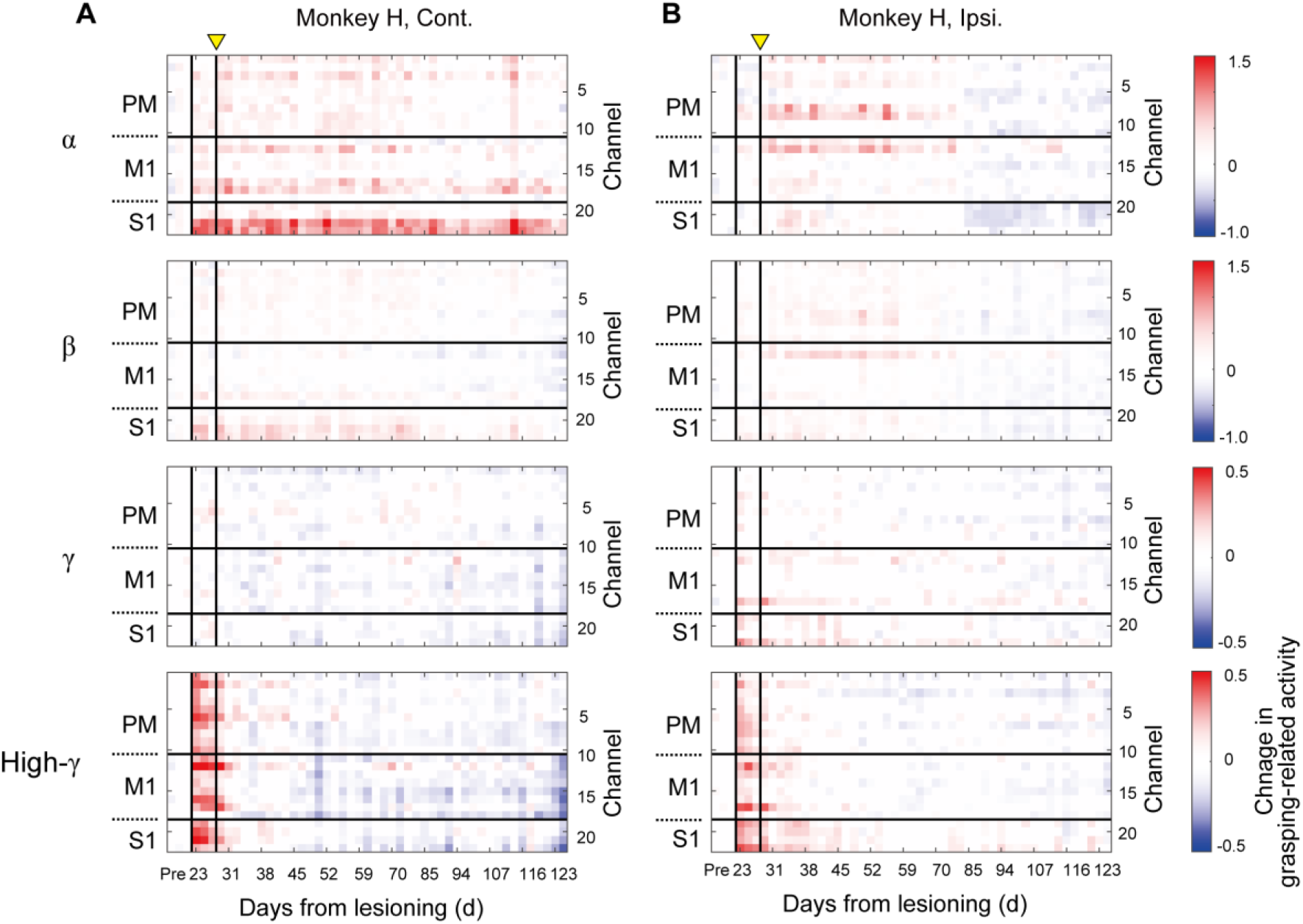
Longitudinal change in peri-movement electrocorticography activity after spinal cord injury shown in Fig. S7, yielded by subtraction of the prelesion data in monkey H. M1, primary motor; PM, premotor; S1, primary somatosensory.

**Fig. S9.**
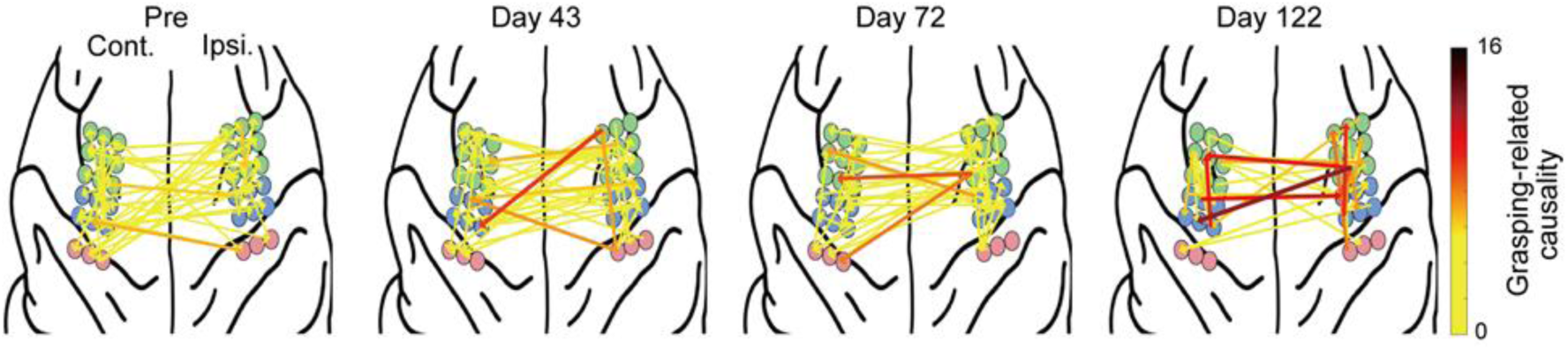
Examples of Granger causality of peri-movement activity in the α band among all combinations of electrocorticography electrodes in the bilateral premotor (PM), primary motor (M1), and primary somatosensory (S1) areas before (Pre) and after (Days 43, 72, and 122) lesioning in monkey M. The top 5% of connections are shown. Note that Granger causality from the ipsilesional PM/M1 to contralesional PM/M1 was enhanced after spinal cord injury.

**Fig. S10.**
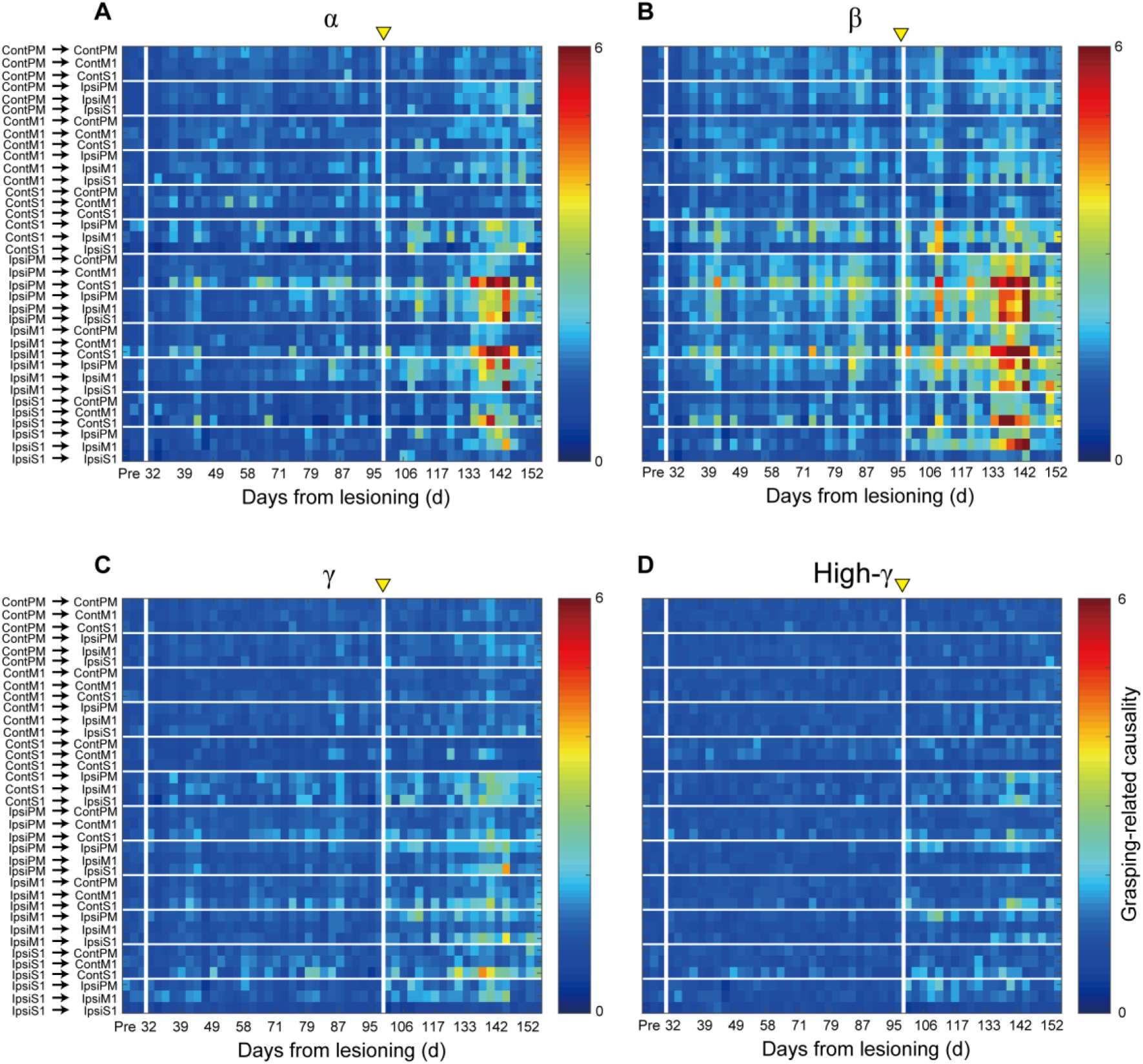
Longitudinal analysis of Granger causality of peri-movement activity at the α, β, γ, and high-γ frequency ranges between the electrocorticography channels in different subregions (contralesional or ipsilesional premotor [PM], primary motor [M1], or primary somatosensory [S1] area) in monkey M. The vertical white line on Day 27 indicates the timing when recovery was saturated (see Fig. 1D–F). Note that α or β band Granger causality from the ipsilesional PM/M1 to contralesional PM/M1 was enhanced around Day 95.

**Fig. S11.**
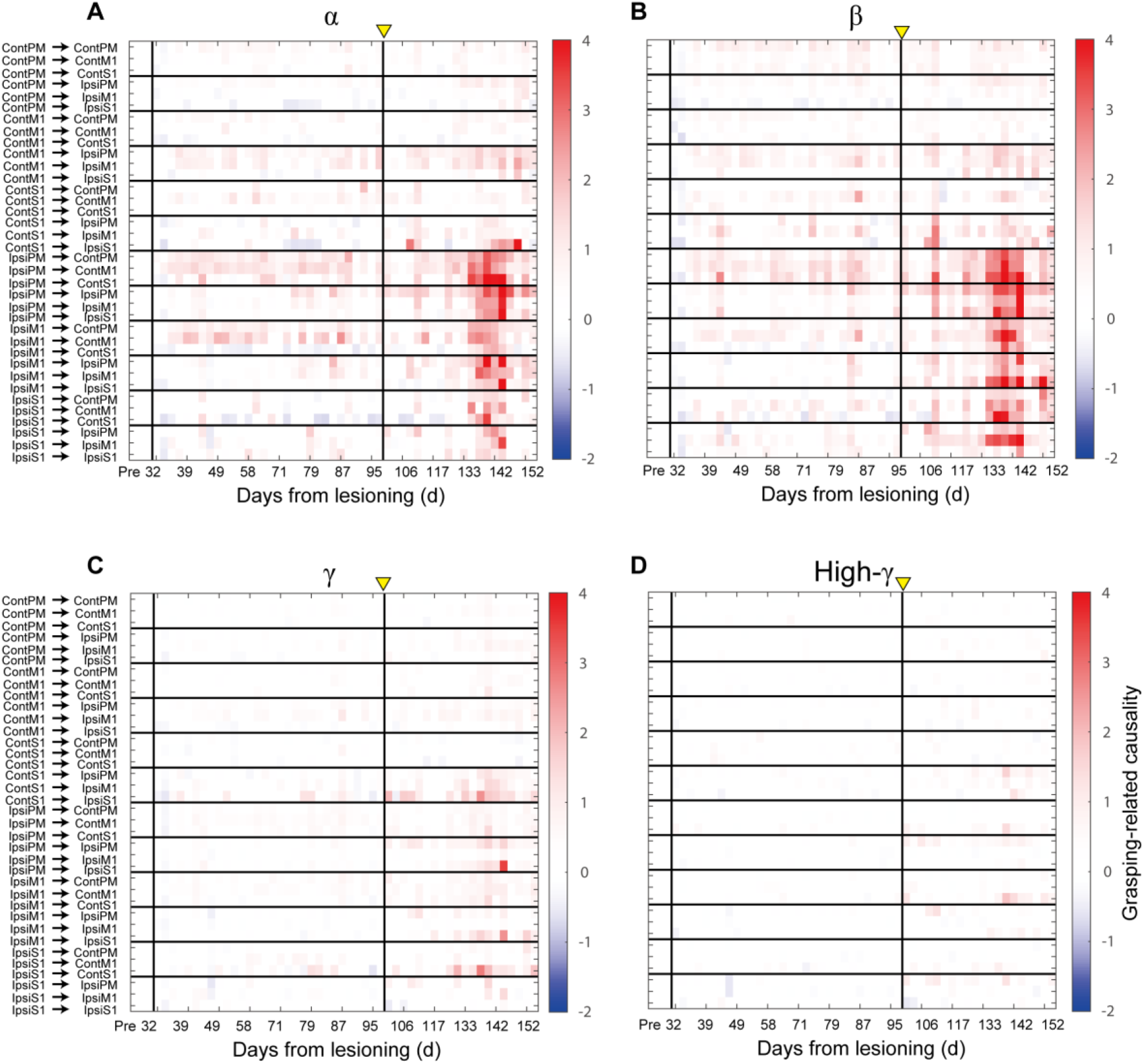
Longitudinal change in Granger causality of peri-movement activity at the α, β, γ, and high-γ frequency ranges between the electrocorticography channels in different subregions (contralesional or ipsilesional premotor [PM], primary motor [M1], or primary somatosensory [S1] area) in monkey M shown in Fig. S10, yielded by subtraction of the prelesion data in monkey M.

**Fig. S12.**
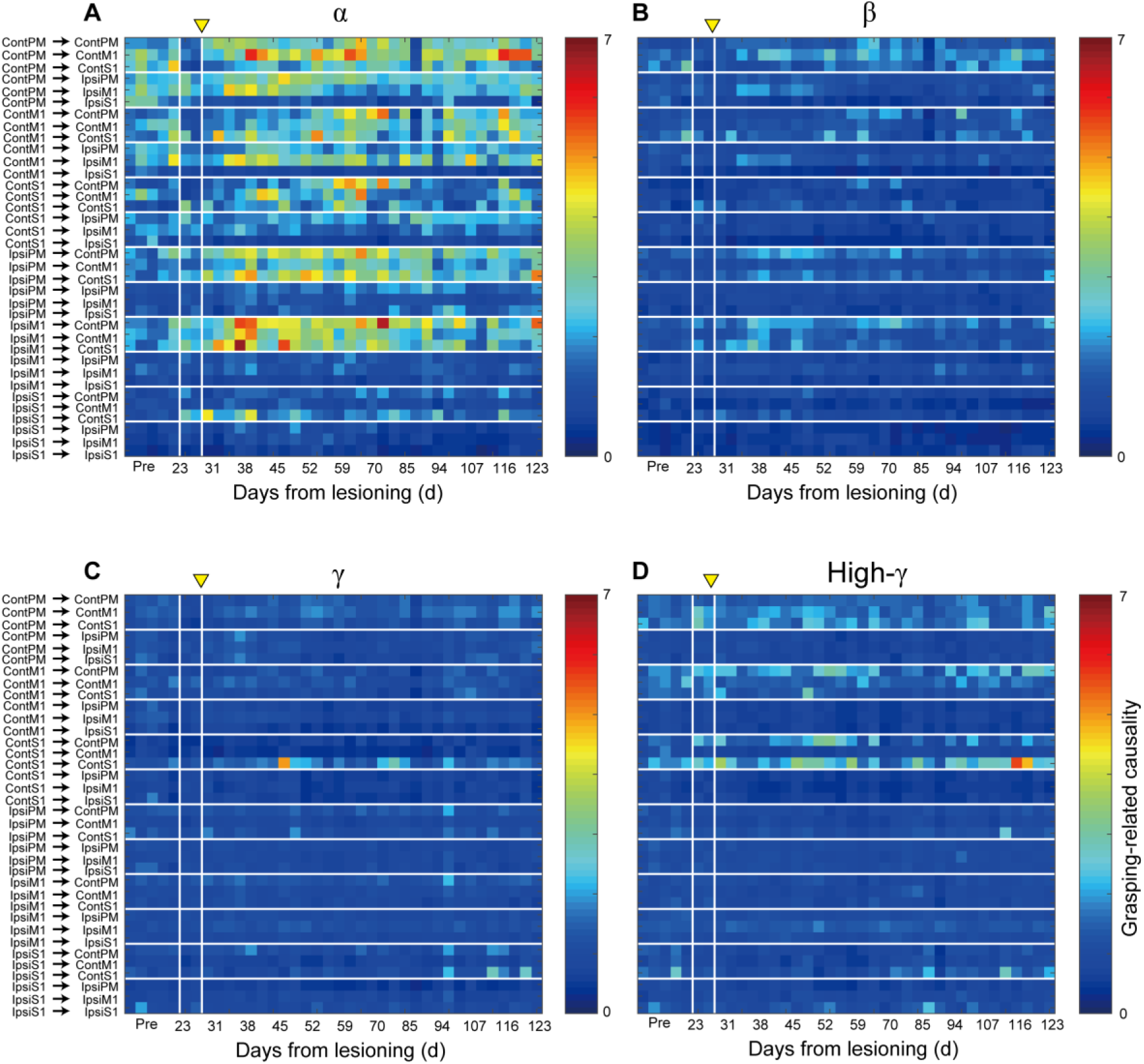
Longitudinal analysis of Granger causality of peri-movement activity at the α, β, γ, and high- γ frequency ranges between the electrocorticography channels in different subregions (contralesional or ipsilesional premotor [PM], primary motor [M1], or primary somatosensory [S1] area) in monkey H. The vertical white line on Day 27 indicates the timing when recovery was saturated (see Fig. 1D– F). Note that α or β band Granger causality from the ipsilesional PM/M1 to contralesional PM/M1 was enhanced after Day 27.

**Fig. S13.**
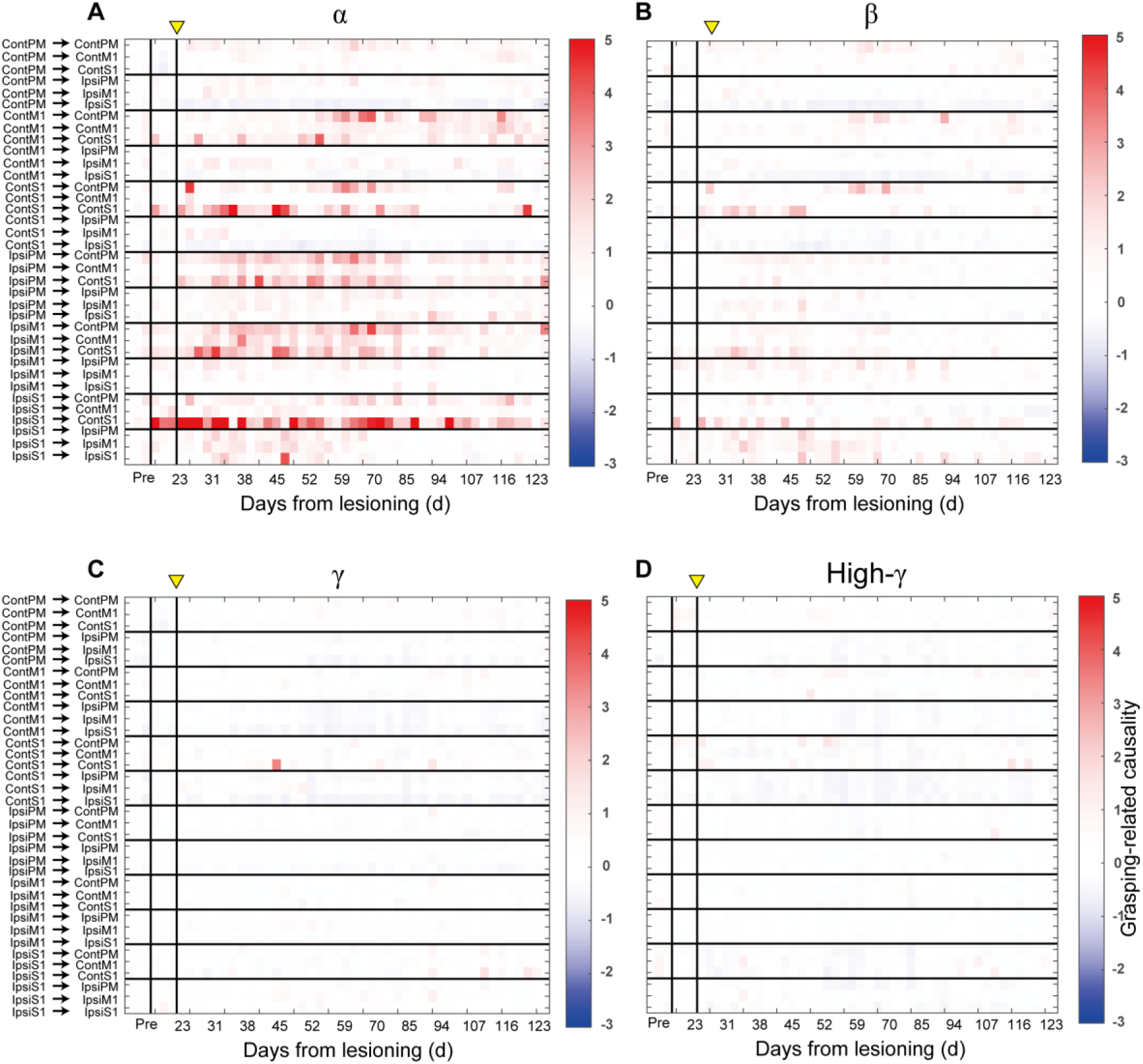
Longitudinal change in Granger causality of peri-movement activity at the α, β, γ, and high- γ frequency ranges between the electrocorticography channels in different subregions (contralesional or ipsilesional premotor [PM], primary motor [M1], or primary somatosensory [S1] area) in monkey H shown in Fig. S12, yielded by subtraction of the prelesion data in monkey H.

**Fig. S14.**
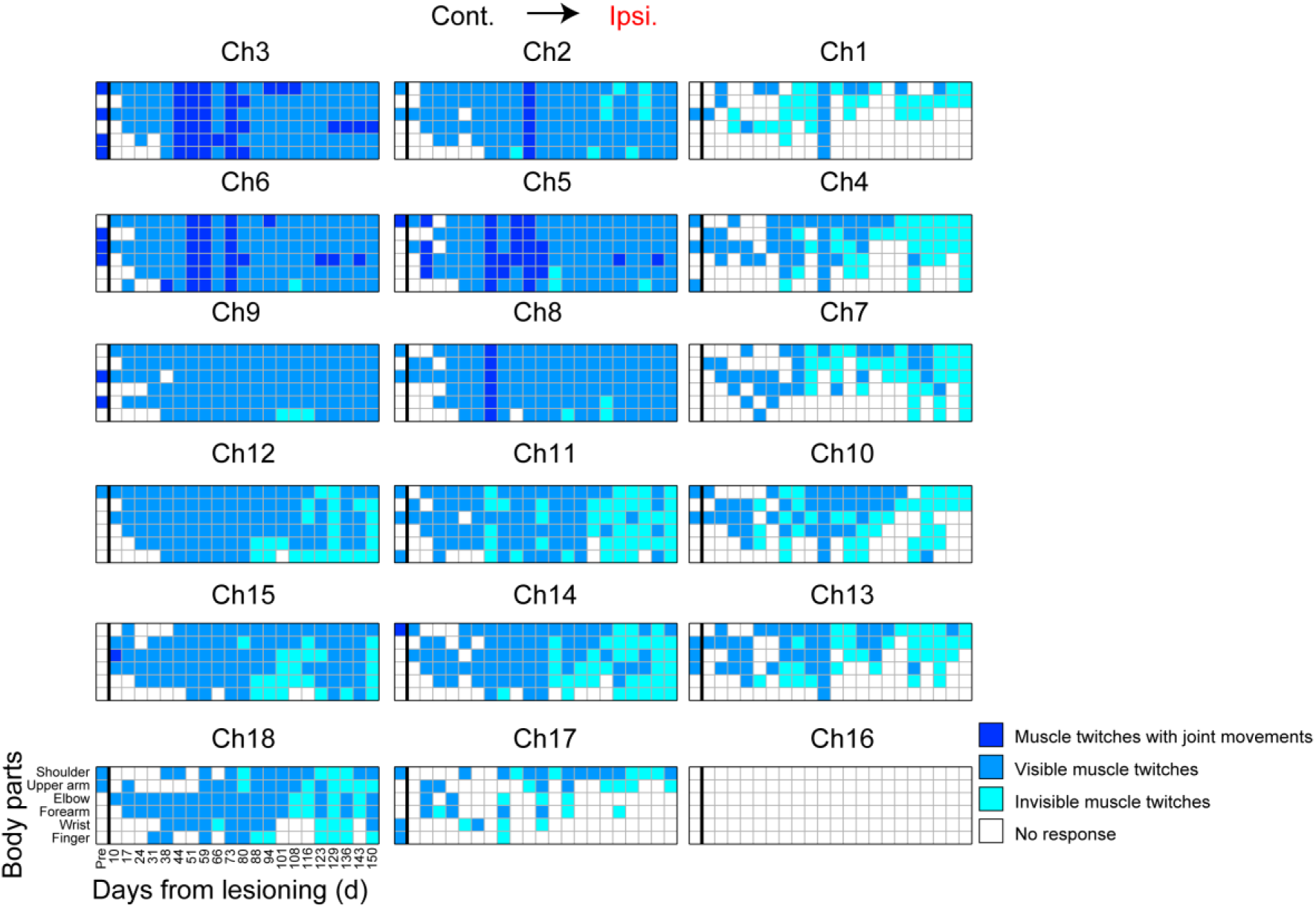
Longitudinal analysis of the magnitude of twitch responses of different body parts (shoulder, upper arm, elbow, forearm, wrist, and finger) on the ipsilesional side induced by stimulation through electrocorticography channels (Ch1–18) on the contralesional premotor, primary motor, and primary somatosensory areas before (Pre) and after (Days 10–150) lesioning in monkey M.

**Fig. S15.**
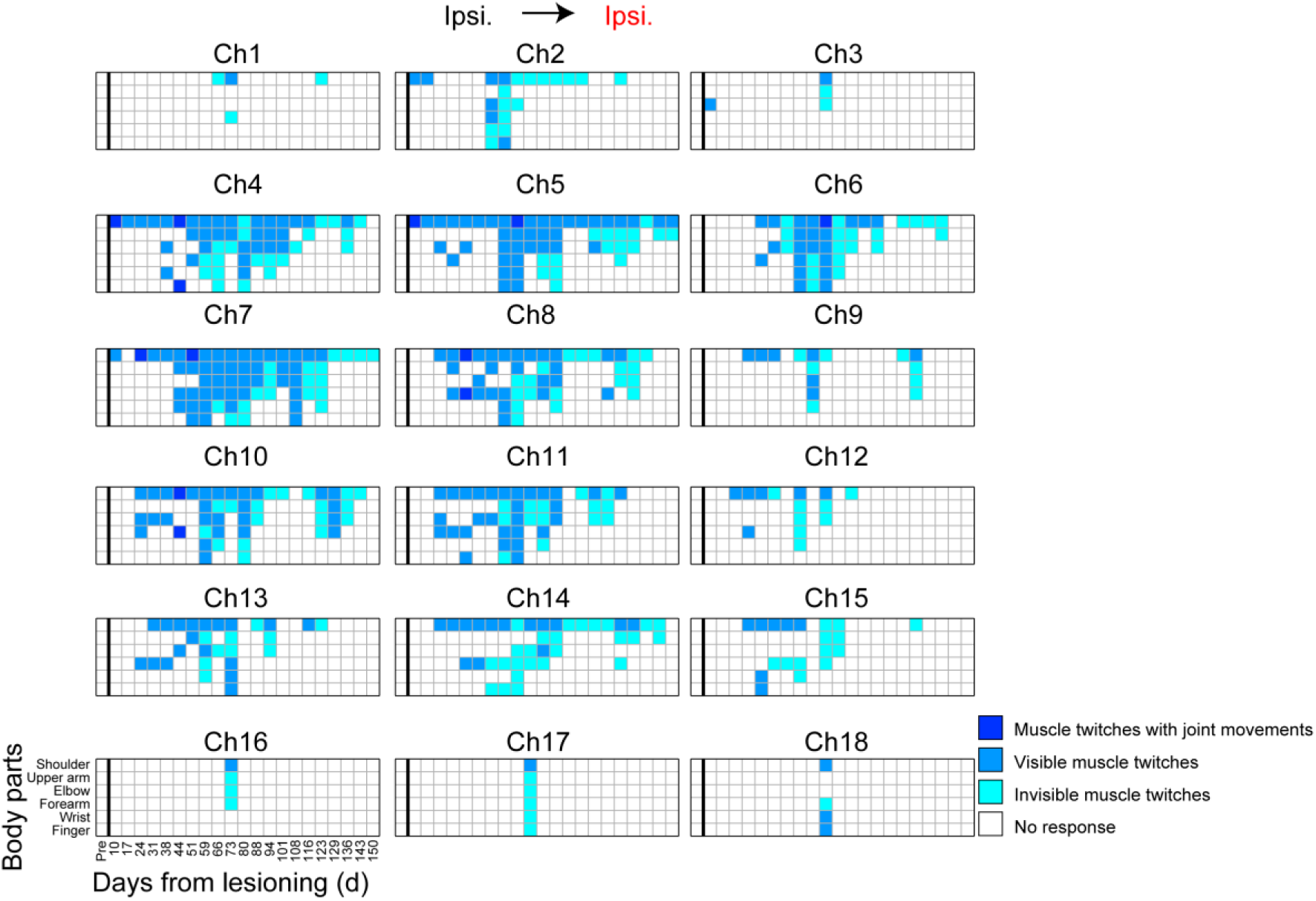
Longitudinal analysis of the magnitude of twitch responses of different body parts (shoulder, upper arm, elbow, forearm, wrist, and finger) on the ipsilesional side induced by stimulation through electrocorticography channels (Ch1–18) on the ipsilesional premotor, primary motor, and primary somatosensory areas before (Pre) and after (Days 10–150) lesioning in monkey M.

**Fig. S16.**
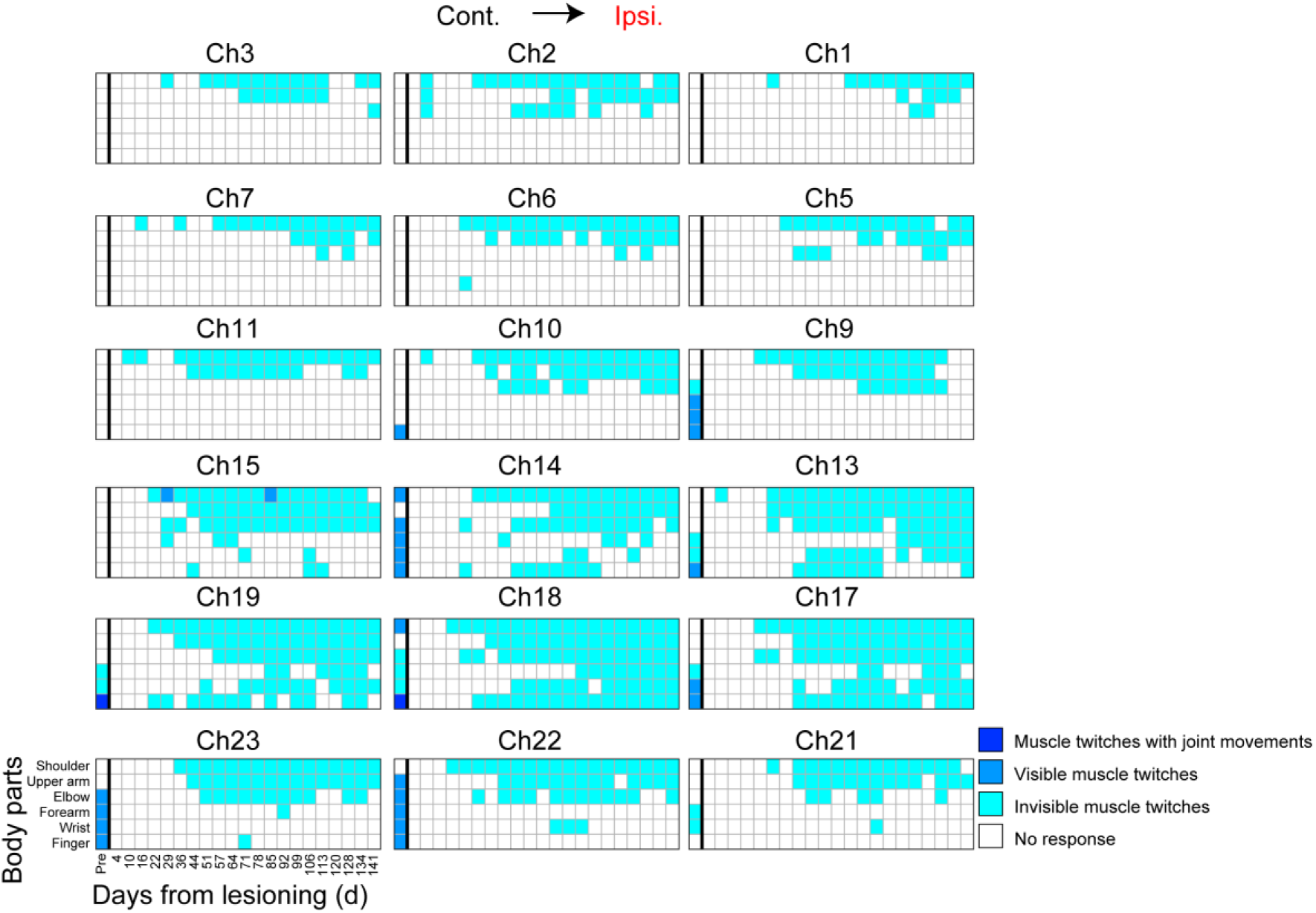
Longitudinal analysis of the magnitude of twitch responses of different body parts (shoulder, upper arm, elbow, forearm, wrist, and finger) on the ipsilesional side induced by stimulation through electrocorticography channels on the contralesional premotor, primary motor, and primary somatosensory areas before (Pre) and after (Days 4–141) lesioning in monkey H.

**Fig. S17.**
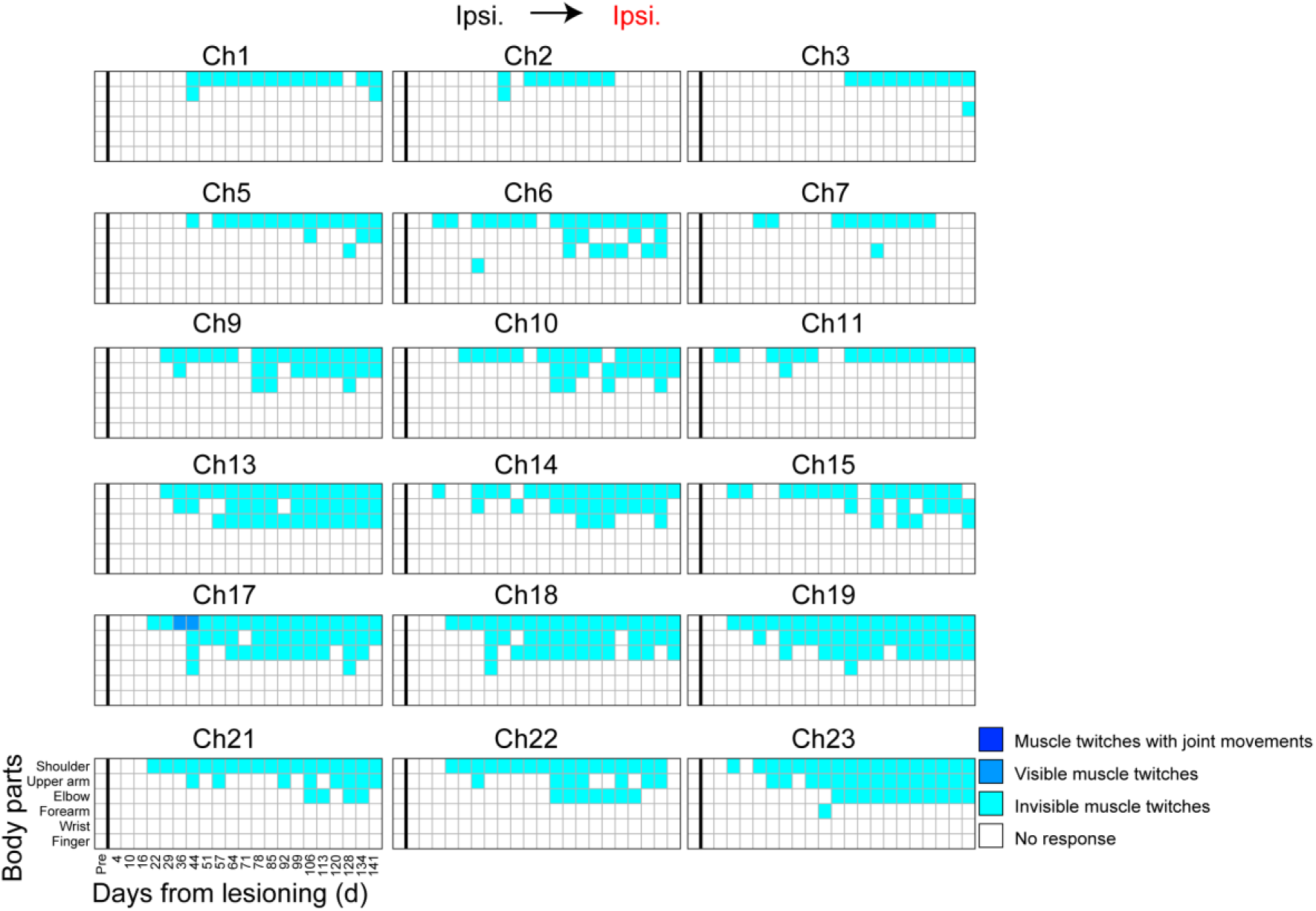
Longitudinal analysis of the magnitude of twitch responses of different body parts (shoulder, upper arm, elbow, forearm, wrist, and finger) on the ipsilesional side induced by stimulation through electrocorticography channels on the ipsilesional premotor, primary motor, and primary somatosensory areas before (Pre) and after (Days 4–141) lesioning in monkey H.

**Fig. S18.**
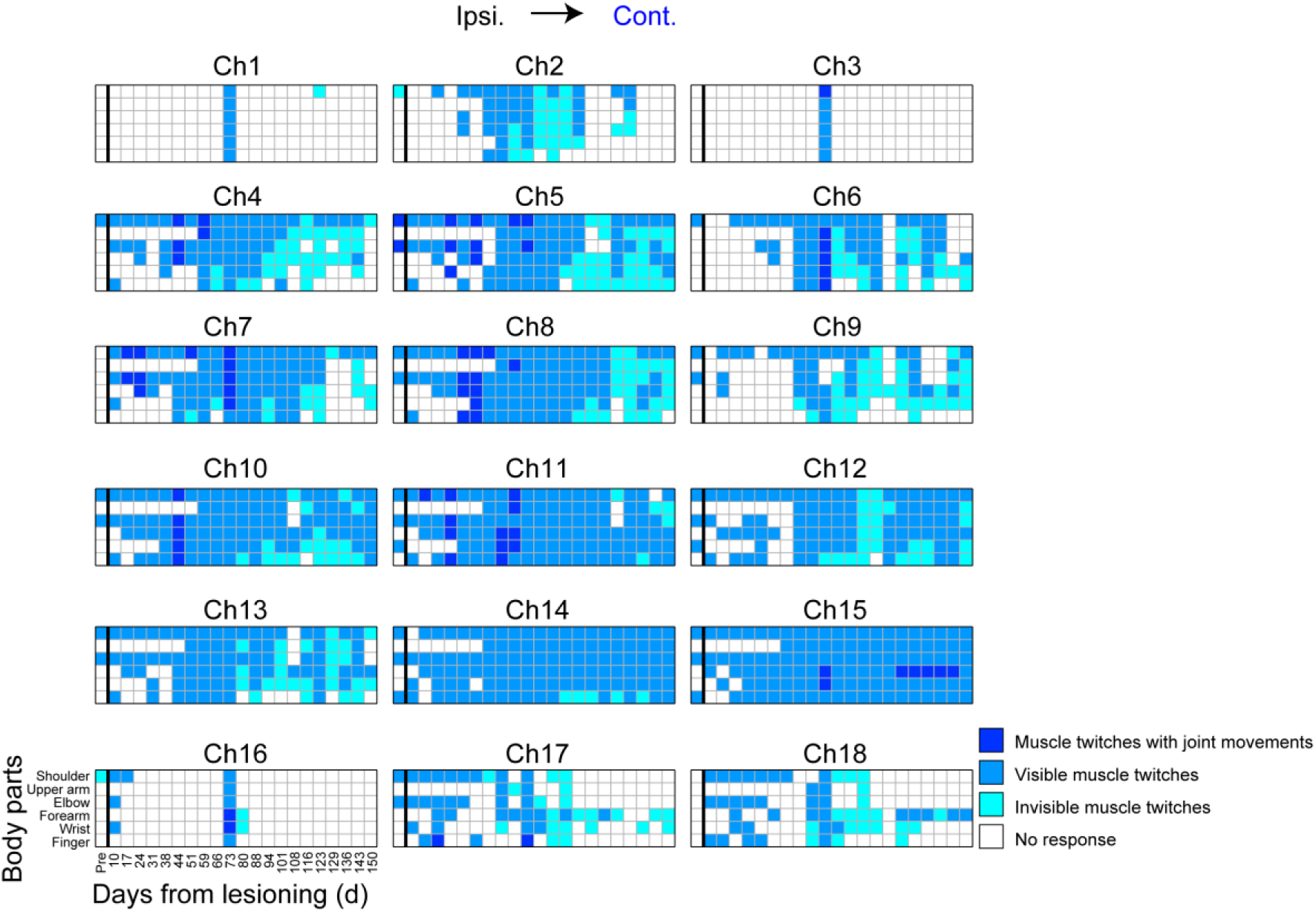
Longitudinal analysis of the magnitude of twitch responses of different body parts (shoulder, upper arm, elbow, forearm, wrist, and finger) on the contralesional side induced by stimulation through electrocorticography channels on the ipsilesional premotor, primary motor, and primary somatosensory areas before (Pre) and after (Days 10–150) lesioning in monkey M.

**Fig. S19.**
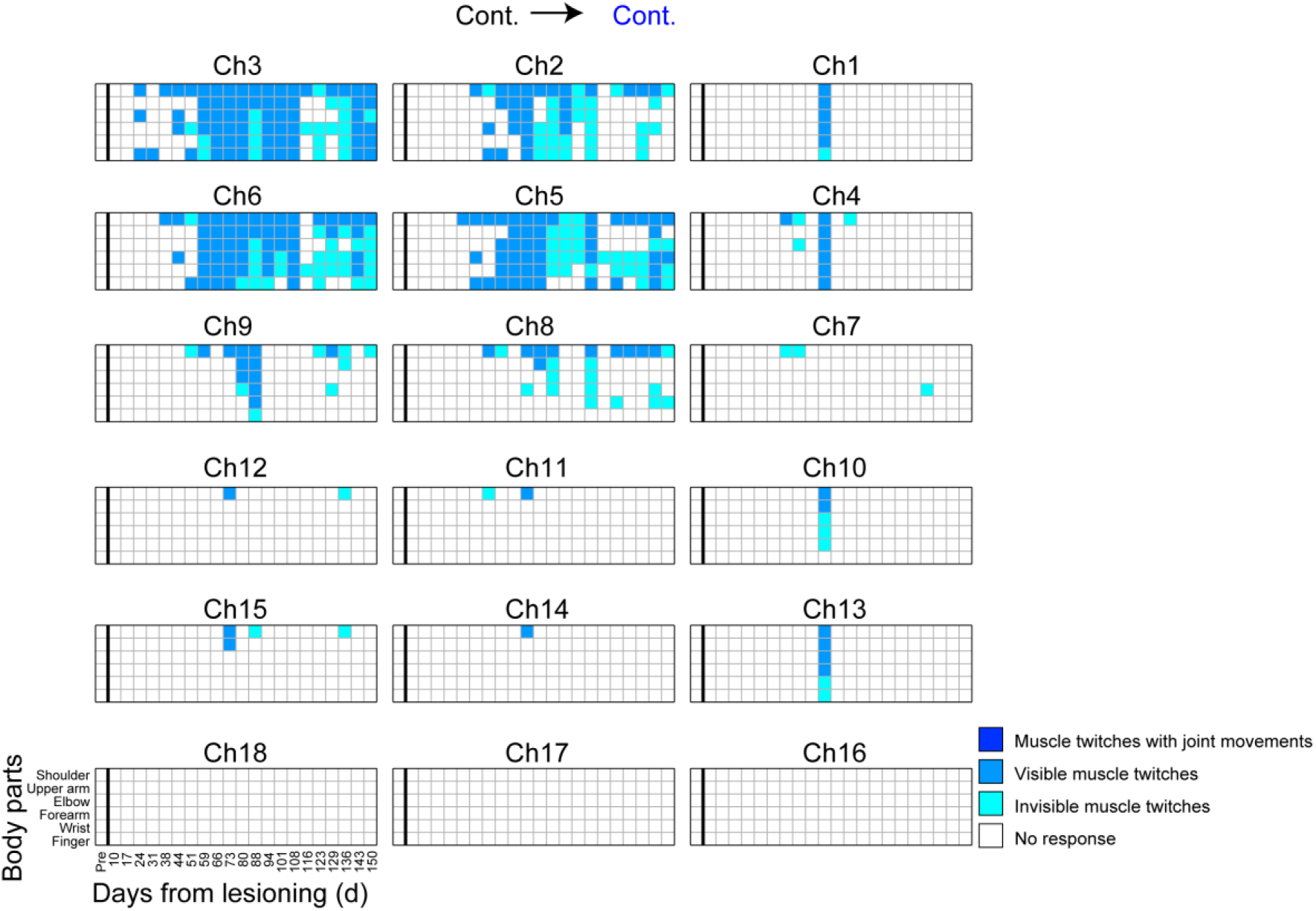
Longitudinal analysis of the magnitude of twitch responses of different body parts (shoulder, upper arm, elbow, forearm, wrist, and finger) on the contralesional side induced by stimulation through electrocorticography channels on the contralesional premotor, primary motor, and primary somatosensory areas before (Pre) and after (Days 10–150) lesioning in monkey M.

**Fig. S20.**
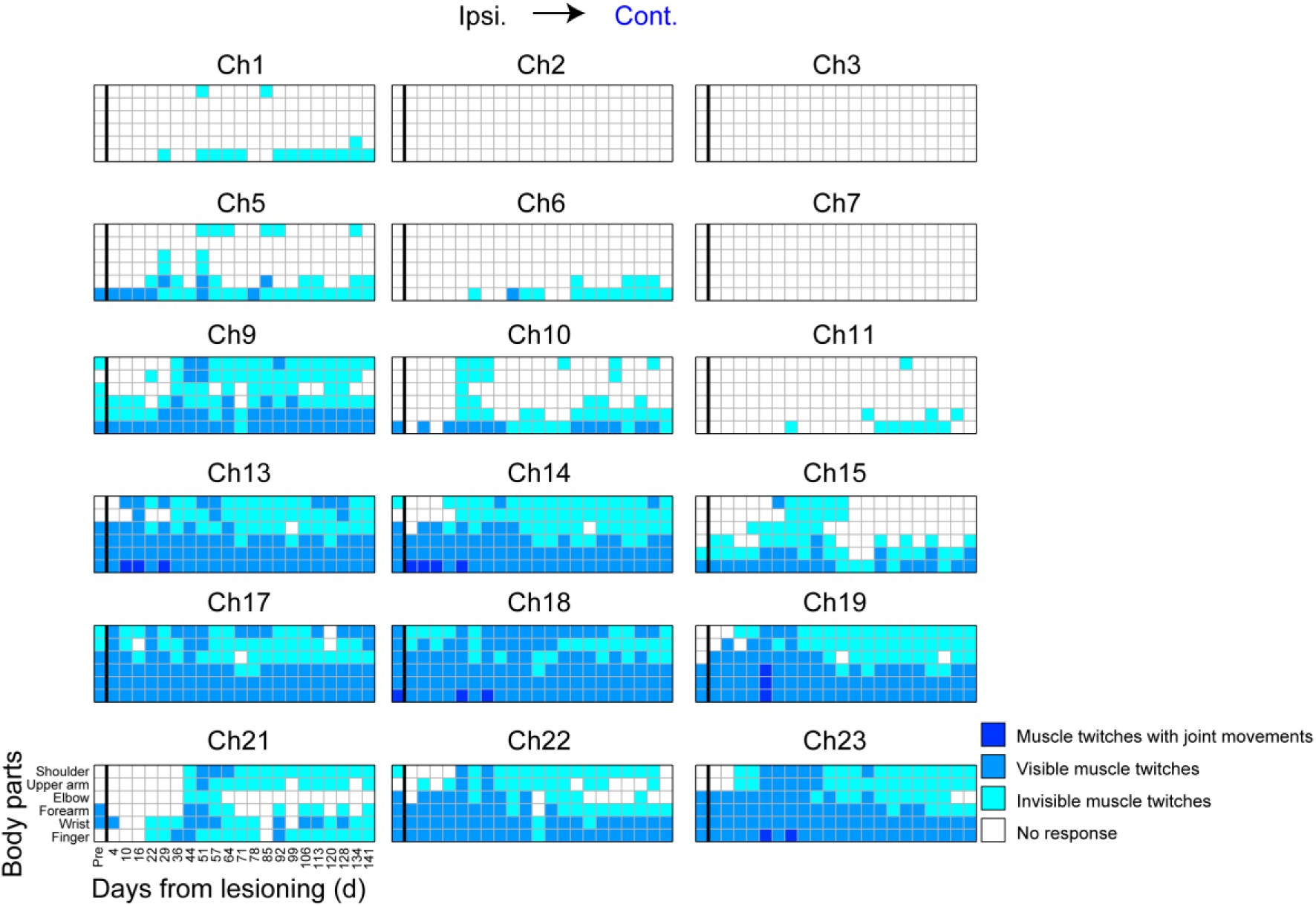
Longitudinal analysis of the magnitude of twitch responses of different body parts (shoulder, upper arm, elbow, forearm, wrist, and finger) on the contralesional side induced by stimulation through electrocorticography channels on the ipsilesional premotor, primary motor, and primary somatosensory areas before (Pre) and after (Days 4–141) lesioning in monkey H.

**Fig. S21.**
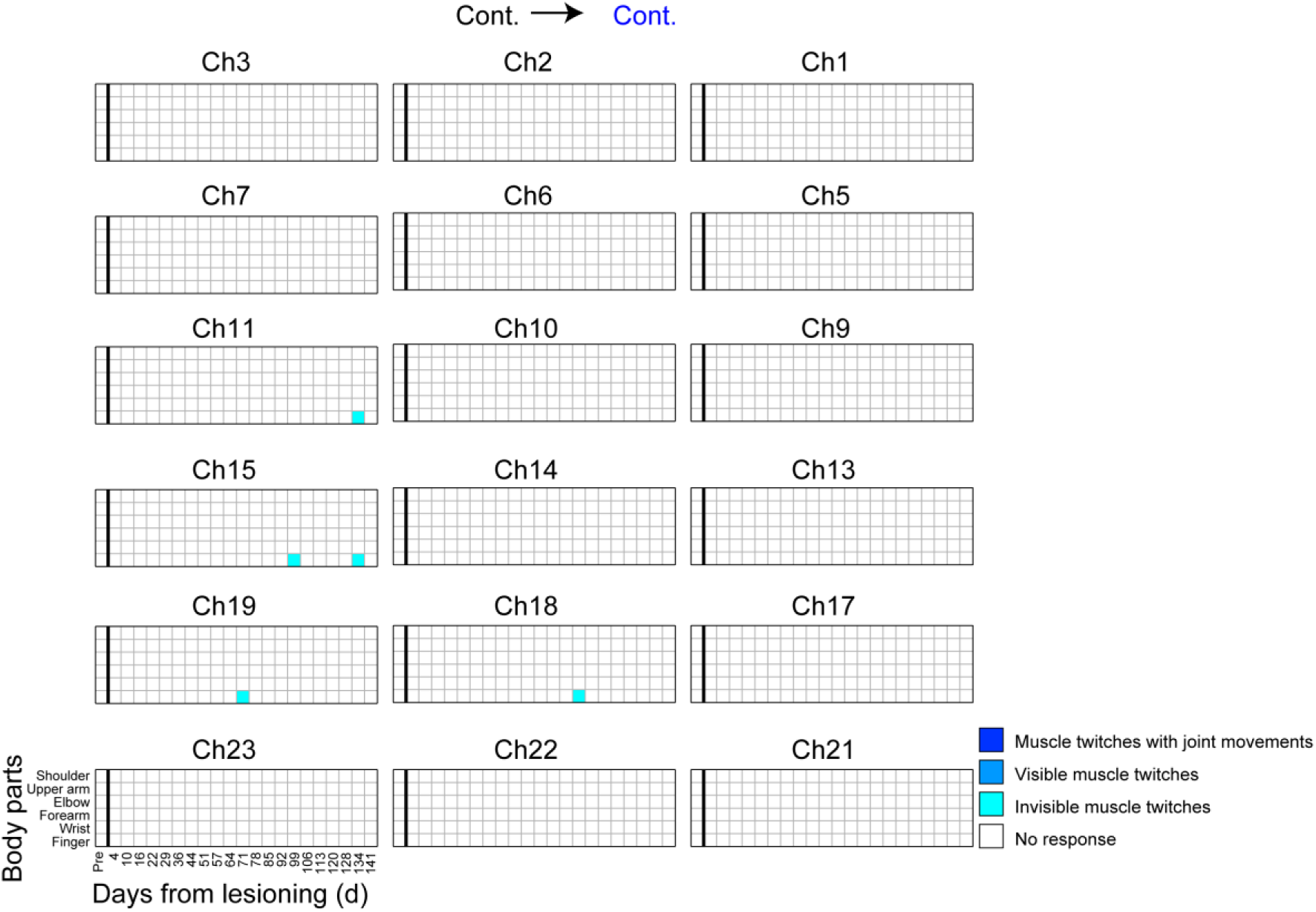
Longitudinal analysis of the magnitude of twitch responses of different body parts (shoulder, upper arm, elbow, forearm, wrist, and finger) on the contralesional side induced by stimulation through electrocorticography channels on the contralesional premotor, primary motor, and primary somatosensory areas before (Pre) and after (Days 4–141) lesioning in monkey H.

**Fig. S22.**
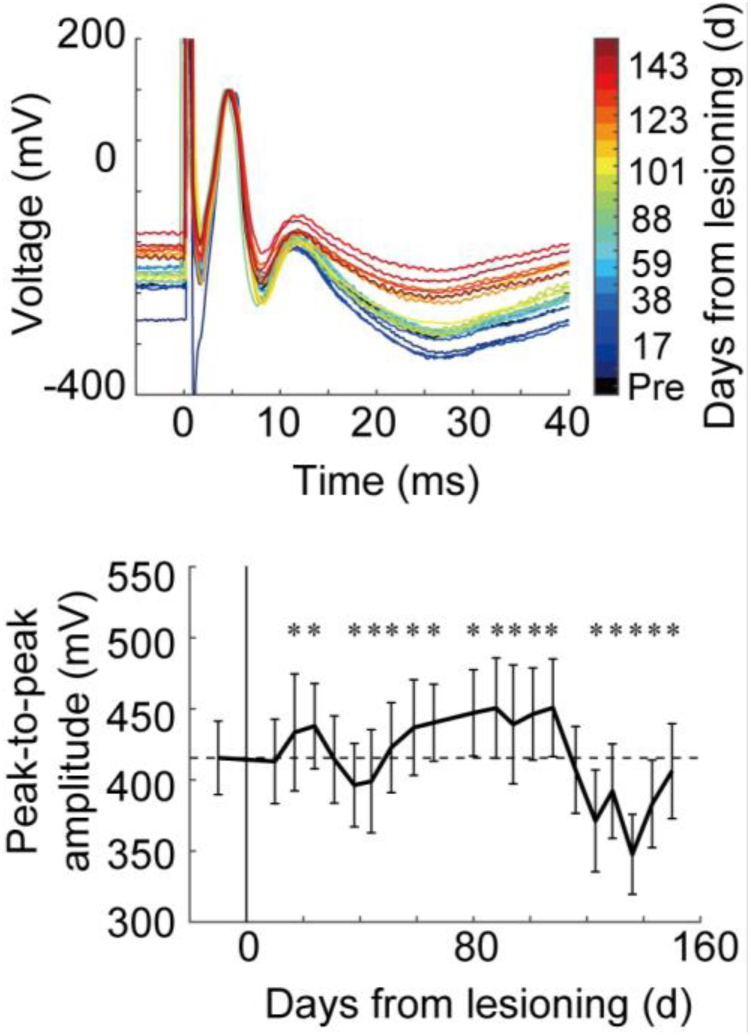
Change in the shape of evoked potentials and peak-to-peak amplitudes. Stimulation was applied to channel (Ch) 8 on the ipsilesional premotor area and recording was from Ch10 on the contralesional primary motor area in monkey M. In the bottom panel, the dotted line indicates the averaged peak-to-peak amplitude in the preoperative stage. The asterisks show a significant difference (p < 0.05, Wilcoxon rank-sum test) in peak-to-peak amplitude between experimental days and preoperative stage.

**Fig. S23.**
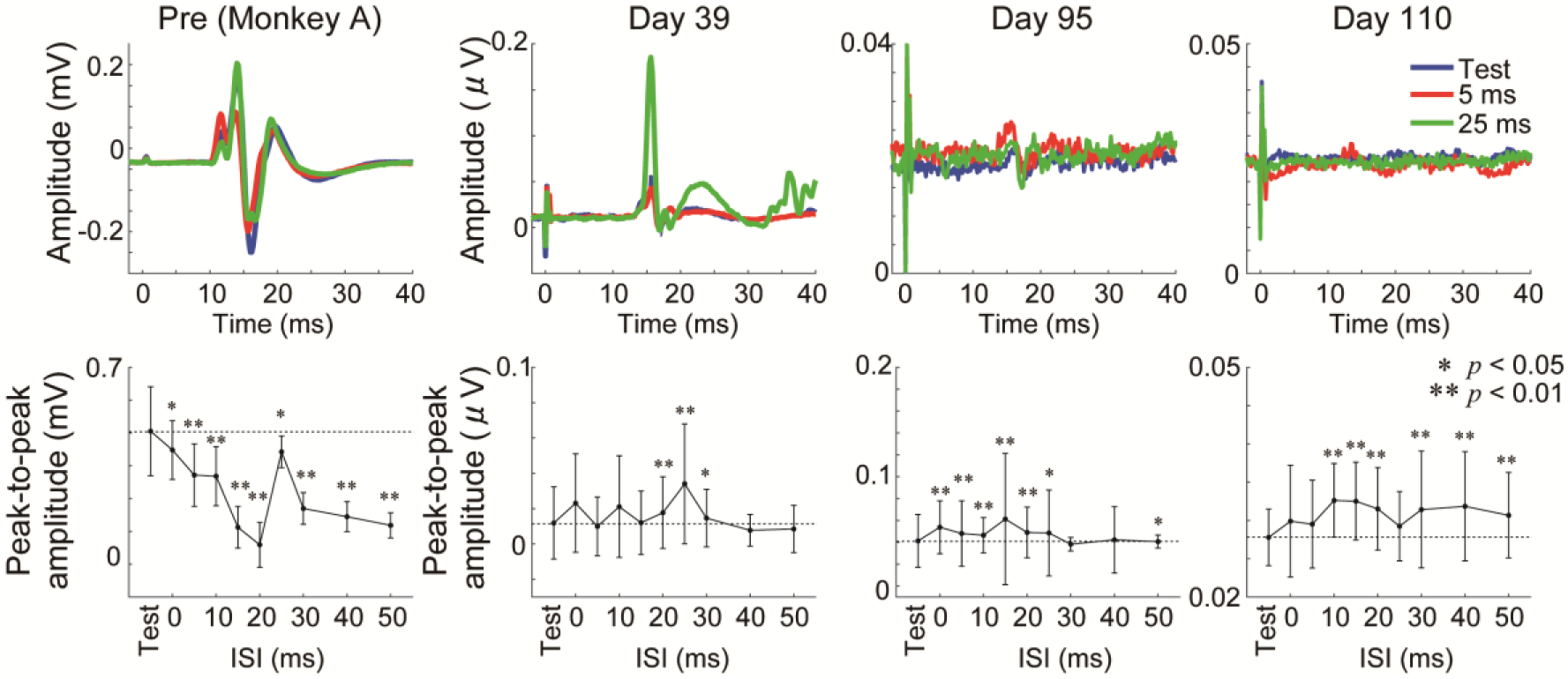
Interhemispheric interactions in monkey A (prelesion) and monkey M (after spinal cord injury). The test stimulus was applied to the contralesional primary motor area (channel [Ch] 11, see Fig. S1) and the conditioning stimulus was given to the ipsilesional premotor area (Ch8, see Fig. S1). The electromyography responses were recorded from the right first dorsal interosseous. Results of the conditioning test before (Pre) and after (Days 39, 95, and 110) lesioning. The upper panels show the superimposed records of the electromyography responses with the test stimulus alone (blue) and with the conditioned stimulus (interstimulus interval [ISI] 5 ms, red; 25 ms, green). The lower panels indicate the averaged peak-to-peak responses with different ISIs. The horizontal dotted line indicates the magnitude of the test response alone. Asterisks indicate a significant difference in peak-to-peak amplitude between the conditioned and test responses (*p < 0.05, **p < 0.01, Wilcoxon rank-sum test).

**Fig. S24.**
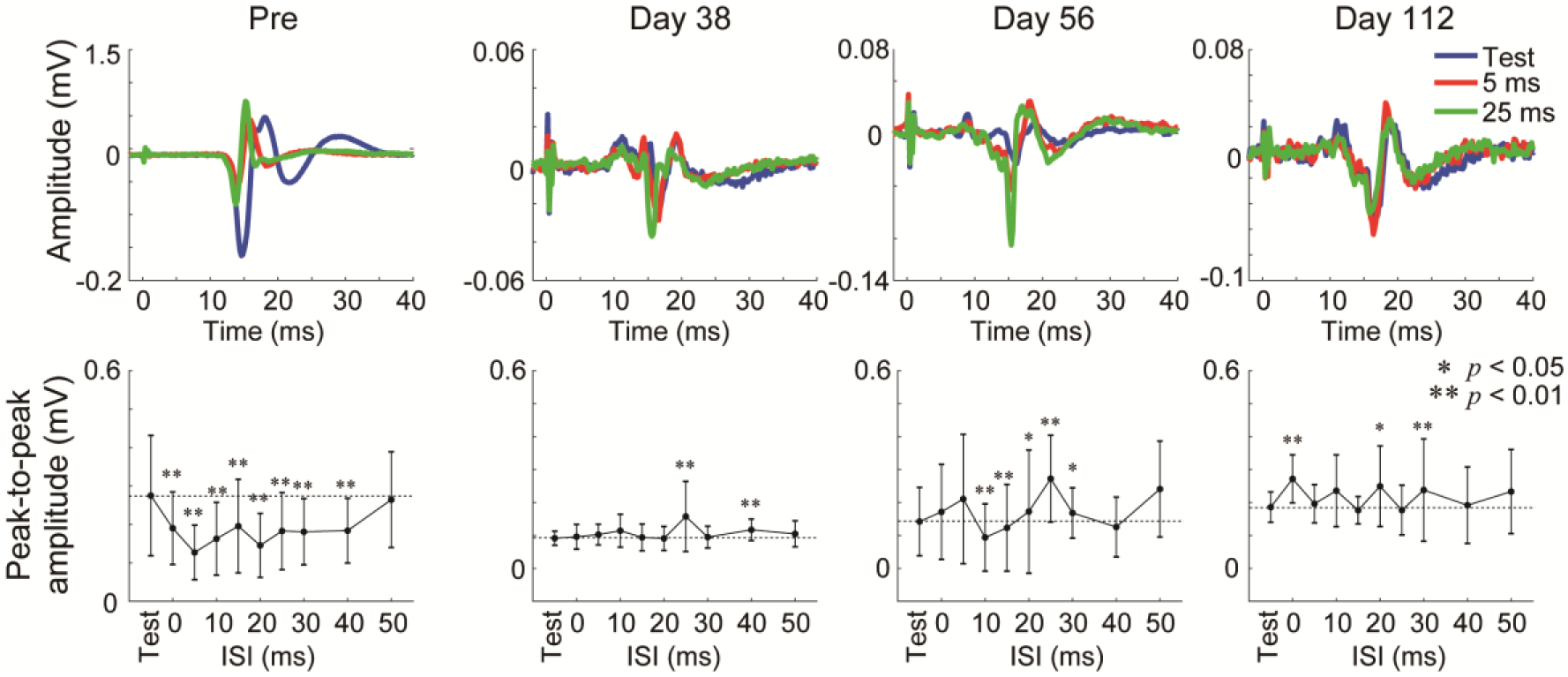
Longitudinal analysis of interhemispheric interactions in monkey H. The test stimulus was applied to the contralesional primary motor area (channel [Ch] 18, see Fig. S1) and the conditioning stimulus was given to the ipsilesional premotor area (Ch11, see Fig. S1). The electromyography responses were recorded from the right first dorsal interosseous. Results of the conditioning test before (Pre) and after (Days 38, 56, and 112) lesioning. The upper panels show the superimposed records of the electromyography responses with the test stimulus alone (blue) and with the conditioned stimulus (interstimulus interval [ISI] 5 ms, red; 25 ms, green). The lower panels indicate the averaged peak-to-peak responses with different ISIs. The horizontal dotted line indicates the magnitude of the test response alone. Asterisks indicate a significant difference in peak-to-peak amplitude between the conditioned and test responses (*p < 0.05, **p < 0.01, Wilcoxon rank-sum test). “Pre” and “Day 56” data are the same as in Fig. 4D.

**Fig. S25.**
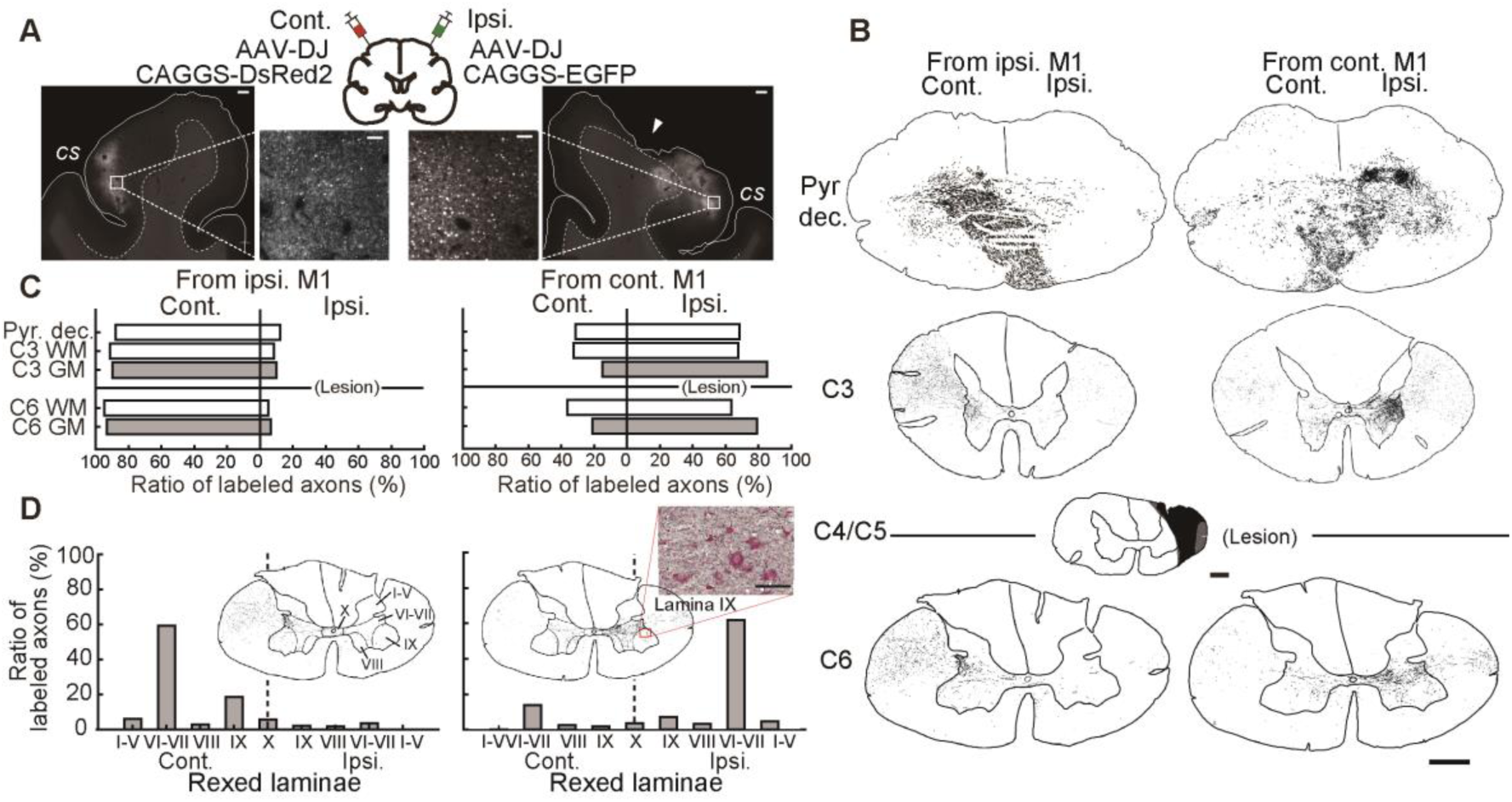
Trajectories of the corticospinal tract after subhemisection in monkey H. (A) Injection sites and labeled cell bodies in the primary motor (M1) area on the contralesional and ipsilesional sides. To account for differences in effect due to the differences of viruses, the vectors were switched between monkeys H and M so that AAV-DJ-CAGGS-DsRed was injected into the contralesional side and AAV-DJ-CAGGS-EGFP was injected into the ipsilesional side. White arrowhead indicates the caving on the surface of the ipsilesional cortex because of the chronic implant of the electrocorticography electrodes, but it did not seem to have caused any serious effects on brain activity (Fig. 2). Scale bars in the lower and higher magnification images are 1,000 µm and 100 µm, respectively. (B) Camera lucida drawings of the labeled axons originating from the ipsilesional (left panels) and contralesional (right panels) M1 at the pyramidal decussation, C3, and C6. Note that at the pyramidal decussation (Pyr. dec.), the axons originating from the ipsilesional M1 (left panel) mainly project from the ipsilesional pyramid to the dorsal part of the contralesional brainstem, while the axons originating from the contralesional M1 (right panel) spread bilaterally to the dorsal brainstem from the pyramid on the contralesional side. Caudal to the lesion (C6), there were some labeled axons from the contralesional M1 on the ipsilesional lateral funiculus at C6. It is considered that these axons were extended after lesioning rather than remaining axons because we confirmed that the lateral funiculus of the spinal cord was completely transected, as shown in Fig. S3, although the lesion size was relatively smaller than in monkey M. Scale bars, 1,000 µm. (C) Proportion of labeled axons in each region of interest (Fig. S26) at each segment. The proportion was calculated by dividing the number of labeled axons on one side by the sum of them on both sides in each segment. The ratio of axons originating from the contralesional side at C6 white matter (WM) leaned to the ipsilesional side because of the presence of putative reconnected axons (see also Fig. S2). Note that the ratio of axons originating from the contralesional M1 leaned to the contralesional side at the pyramidal decussation and recovered to the ipsilesional side at C6 gray matter (GM) where they should have been distributed originally, which is different from the distribution of axons originating from the ipsilesional M1. (D) The laminar distribution of labeled axons in C6 GM in the Rexed laminae (I–V, VI–VII, VIII, IX, and X). The number of labeled axons in each lamina was divided by sum of the axons in the GM. The high magnification image shows that the labeled axons originating from the contralesional M1 project into lamina IX on the ipsilesional side, where they contacted the counterstained presumed motoneurons. Scale bar: 100 µm.

**Fig. S26.**
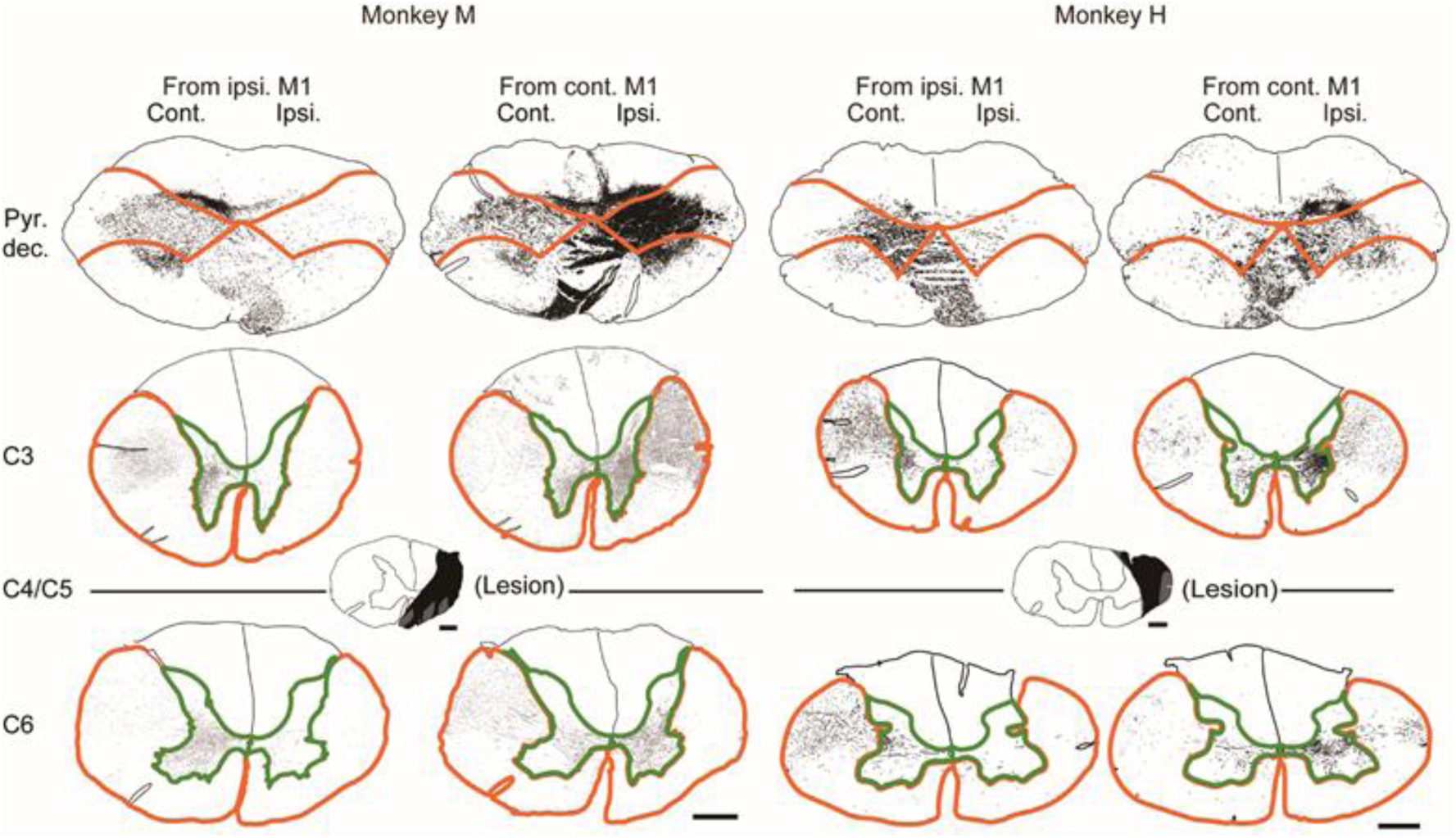
Regions of interest (ROIs) for quantifying the labeled axons (see also Figs. 5C and S25C). The ROIs were determined based on the counterstaining of DAB-stained sections and overlaid to the traced illustration to count the number of labeled axons. The ROIs for the pyramidal decussation (Pyr. dec.) are surrounded by orange lines on the contralesional (Cont.) and ipsilesional (Ipsi.) sides where the labeled axons descending to the caudal segments reside. The medial part with axons projecting to each side densely was mixed, while the dorsal part, which was judged as the cuneate nucleus based on the counterstaining, was excluded. The ROIs for the C3 and C6 segments were defined by the lateral and ventral funiculi outlined with orange lines, which was shown as white matter in Figs. 5C and S25C. The ROIs for gray matter are outlined by green lines. Images are the same as those shown in Figs. 5B and S25B. Scale bars: 1,000 µm. M1, primary motor area.

**Fig. S27.**
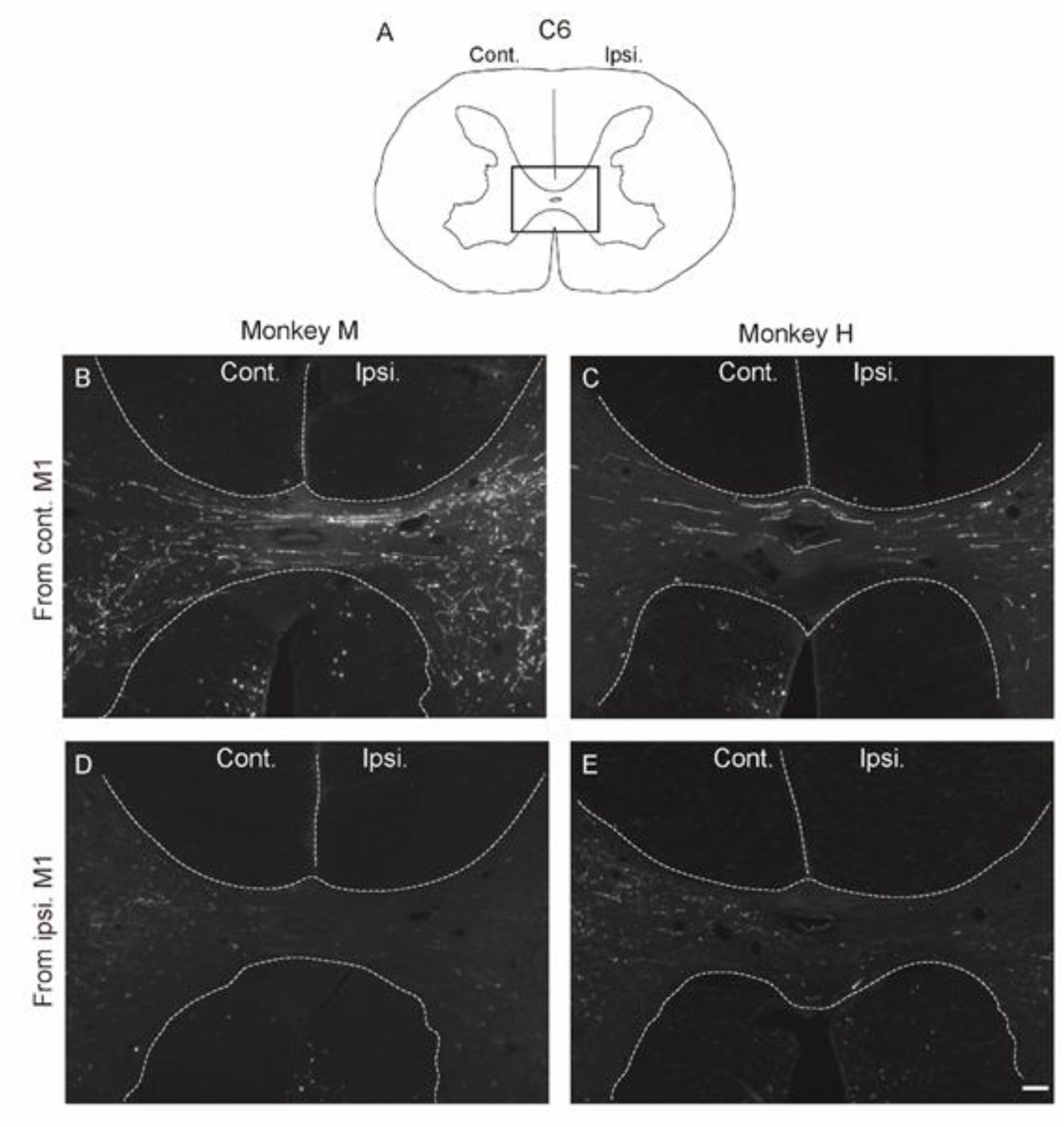
Dark-field images of labeled corticospinal axons with immunofluorescent staining in the medial part of the spinal segments caudal to the lesion. (A) The areas of the photomicrographs in (B– E) at the C6 transverse section (square). (B, C) Distribution of the labeled axons originating from the contralesional primary motor (M1) area in monkeys M (left panel) and H (right panel). (D, E) Distribution of the labeled axons originating from the ipsilesional M1 in monkeys M (left panel) and H (right panel). Scale bar: 100 µm. Note that many axons originating from the contralesional M1 crossed over the midline and were distributed on the ipsilesional side of the gray matter in both monkeys, whereas almost no axons originating from the ipsilesional side crossed over the midline. These results suggested that the axons originating from the contralesional M1 crossed the midline from left to right, that is, were re-routed to the ipsilesional side caudal to the lesion.

**Table S1.**
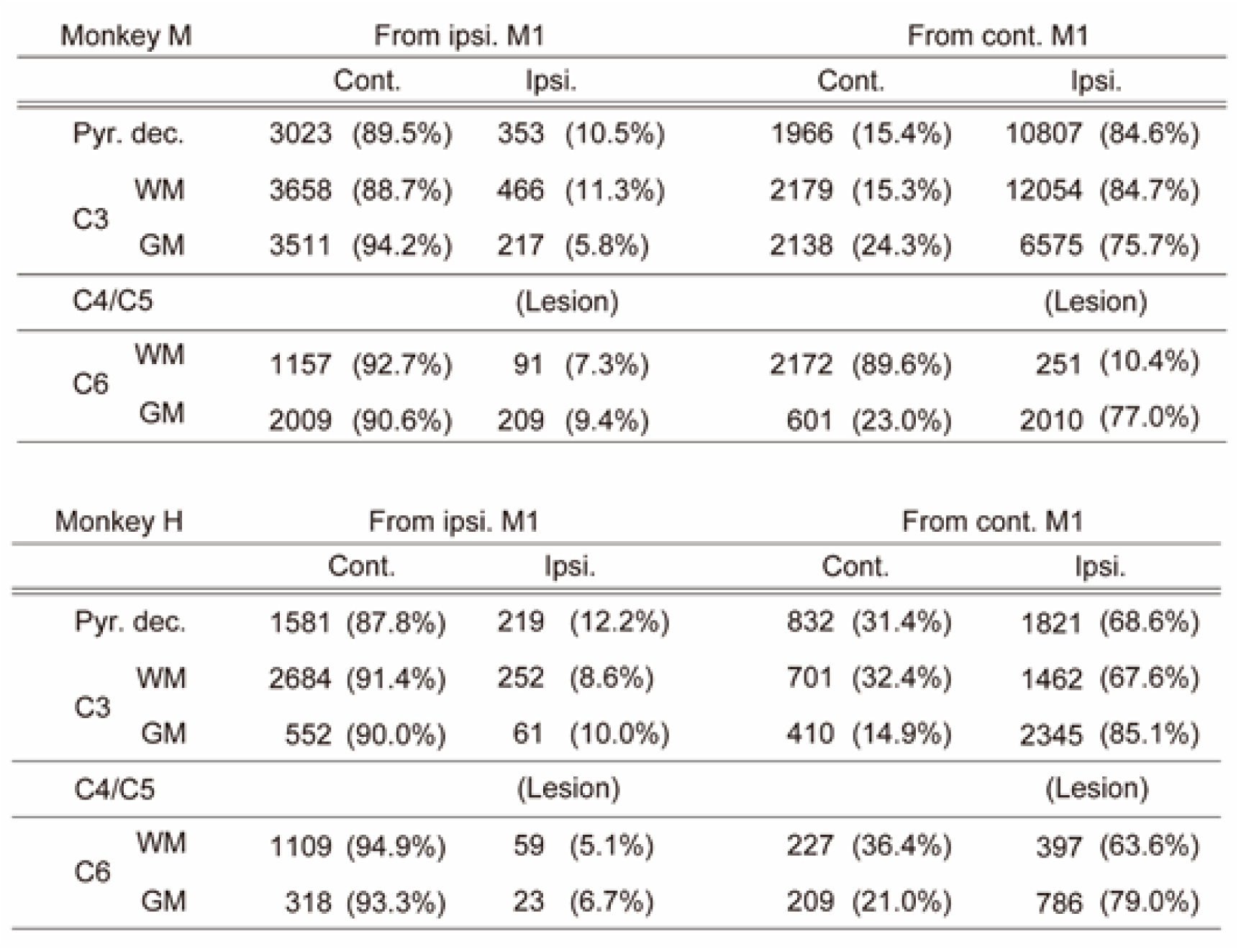
Absolute number of traced axons in each section at the level of the pyramidal decussation (Pyr. dec.), C3, and C6 in each monkey (see also Figs. 5C and S25C). For the axons originating from the ipsilesional (Ipsi.) primary motor (M1) area, approximately 90% were counted on the contralesional side (Cont.) and 10% were on the ipsilesional side through the spinal cord in both animals, as reported in intact monkeys (17, 18). Note that the proportion of labeled axons originating from the contralesional M1 on the ipsilesional side was much higher than 10% at the pyramidal decussation and this trend was continued to the white matter (WM) at C3. Furthermore, at C6, despite the lesion being just rostral to this level, as shown in Figs. S2 and S3, almost 80% of the labeled axons were detected in the ipsilesional gray matter (GM) in both animals.

Movie S1. Recovery process in monkey M

Movie S2. Recovery process in monkey H

## Notes

### Competing Interest Statement

The authors have declared no competing interest.

